# Hollow-fibre biomanufacturing and cell-free engineering of HEK293 extracellular vesicles

**DOI:** 10.1101/2025.09.20.677537

**Authors:** Richard J. R. Kelwick, Alexander J. Webb, Amelie Heliot, Paul S. Freemont

**Author notes:** Joint first authors (Richard J. R. Kelwick, Alexander J. Webb). **Correspondence to:** Prof. Paul Freemont, Department of Infectious Disease, Imperial College London, London, SW72AZ, UK.

## Abstract

Extracellular vesicles (EVs) are lipid-delineated nanoparticles that are produced by most cell types. EVs contain complex molecular cargoes that can have useful therapeutic or vaccine immunological effects. Cell-free gene expression systems can be used to produce membrane proteins *in vitro*, that can co-localise and integrate with exogenously added EVs. To advance this type of cell-free EV engineering we manufactured, isolated and characterised HEK293 cell EVs. These EVs were successfully cell-free engineered with several CD63-based membrane fusion proteins. In our most optimal conditions, up to 4.83 ×10^11^ /ml of HEK293 EVs were successfully cell-free engineered with a fusion membrane protein incorporating CD63 I-shaped membrane-insertion topology transmembrane helix 3 (CD63ITM3) and monomeric green lantern (mGL). Finally, we also demonstrated that nano flow cytometry is a powerful tool for assessing cell-free EV engineering efficiency. In the future cell-free EV engineering could help accelerate future EV discoveries and the development of EV translational applications.

**Graphical abstract:** 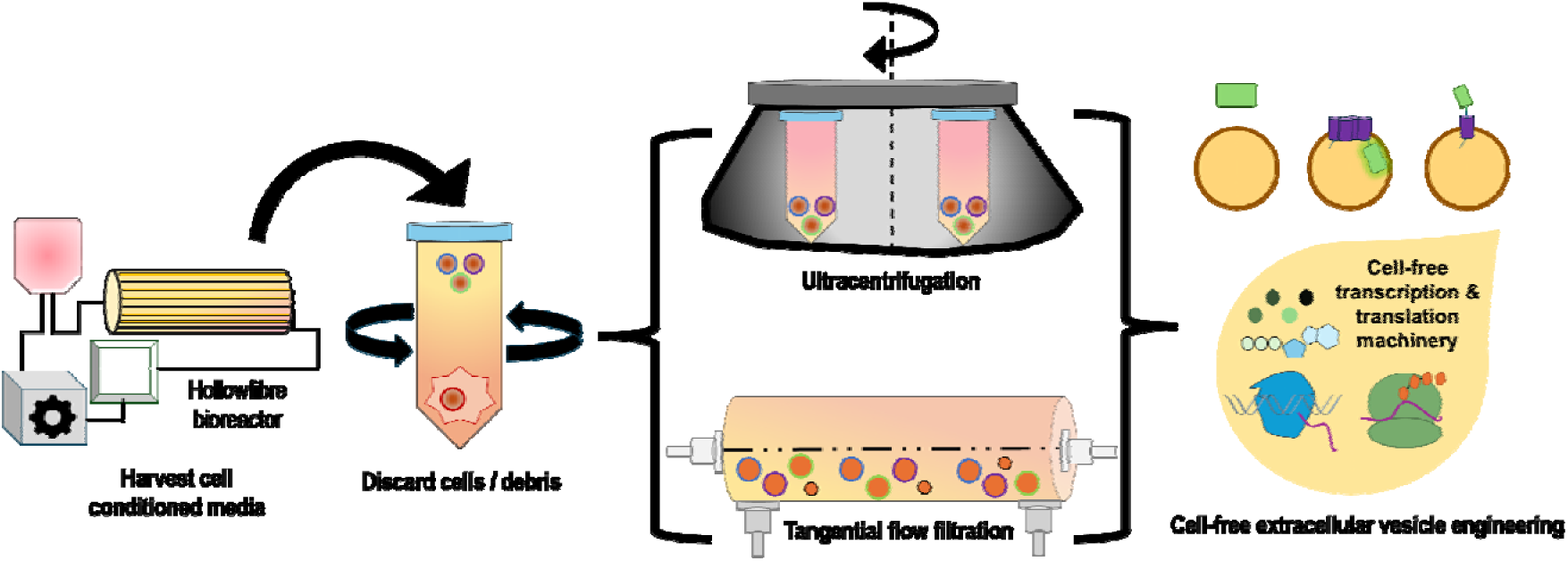

**Technology readiness:** Cell-free systems are a well-established part of the engineering biology toolkit. Indeed, several applications are already at TRL9 stage, including commercially available cell-free protein synthesis kits, drug screening assays, field-tested medical biosensors, as well as therapeutic antibody manufacturing that has now reached cell-free reaction volume scales of up to 4500 L. However, cell-free extracellular vesicle (EV) engineering is currently at TRL3 / 4 stage. Whilst our study helps to further advance cell-free EV engineering, key challenges remain. To accelerate technology readiness, improvements in scalable EV isolation methods, increased eukaryotic cell-free protein synthesis yields, and assay automation will likely speed up cell-free EV engineering workflows. Furthermore, artificial intelligence-guided design workflows could be used to rapidly create community accessible libraries of EV membrane protein scaffolds, therapeutic, cell targeting and other modular elements for use in EV engineering studies. Finally, the development of EV potency assays for functional assessment of cell-free engineered EVs are needed to progress prototyped designs towards clinical translation. If these challenges are addressed, we envision that cell-free EV engineering will become an important approach in EV biological research and clinical applications.

**Highlights:**

- Cell-free extracellular vesicle (EV) engineering could be utilised to rapidly prototype and test novel biotechnological, vaccine, or therapeutic EVs for foundational or translational applications.
- Our data highlight some of the impacts that hollow fibre-based cell culture, EV isolation methods and different cell-free engineering approaches have on cell-free EV engineering workflows.
- We demonstrate several characterisation assays and technologies, including nanoflow cytometry, that can be used to assess cell-free EV engineering efficiency.

## Introduction

Extracellular vesicles (EVs) are lipid-delineated nanoparticles produced by most cell types that have important biological roles [1,2]. EVs also hold significant potential for use in disease diagnostic and therapeutic applications [3–5]. Furthermore, EV molecular cargoes and physiological functions can differ according to cell type of origin, the biogenesis mechanisms by which EVs are shed or secreted and changes in cellular states during health and disease [1,2,6,7]. These contexts can greatly alter EV heterogeneity in terms of the types of EVs that are produced and their cargoes, which can include many different metabolites, nucleic acids, luminal or membrane proteins [6,7]. The biological importance of EVs, along with their accessibility from all biofluids including blood, urine, saliva and cerebral spinal fluid has accelerated interest in the study of EVs as a source of chronic and infectious disease biomarkers [8–10]. Likewise, EV-based vaccines, therapeutics, drug delivery and gene editing vehicles are also highly promising [3,11,12]. Indeed, EVs have several beneficial characteristics including their biocompatibility, ability to deliver complex bioactive molecules to cells and a capacity to cross the blood-brain barrier [13–15]. These EV characteristics could be utilised to treat a wide array of diseases including cancers, inflammatory or neurological conditions and infectious diseases. Arguably, bacterial vesicle-based vaccines including VA-MENGOC-BC and Bexsero are currently the most clinically and commercially successful EVs on the market [16–18]. Though this may change in the future, given that many different therapeutic EVs are currently in pre-clinical and clinical development [3,19]. Interestingly, these more recent studies are expanding beyond bacterial vesicles and are increasingly focusing on human cell-derived EVs, including those from human embryonic kidney (HEK293) cells [20] or mesenchymal stem cells (MSCs) [21,22], and their potential for more sophisticated EV-based therapeutic applications [4,19,23].

Whilst promising, the development of next generation EV therapeutics is not trivial, and many challenges remain. Translational EV research is not only hindered by foundational knowledge gaps in the fields understanding of the heterogeneity of EVs across different biological contexts (*in vivo*), but also in industrial contexts across different cell culture scales (e.g., EVs from bioreactor cultured cells) [2,6,21,24]. This also includes the fields still evolving understanding of the mechanisms by which specific molecules are loaded into or onto EVs [7,15,25]. Another ongoing challenge is that once administered, therapeutic EVs are seemingly quickly removed from circulation or are prone to accumulating in the spleen, lungs or liver, which might impact their therapeutic efficacy and require the use of higher therapeutic EV doses, thereby also increasing adverse drug reaction risks in future patients [26,27]. Finally, upon reaching their cellular targets, the mechanisms by which EVs transfer their cargoes or otherwise influence cells, also requires further investigation [6,15]. Consequently, a deeper understanding of EV biodistribution and the development of novel EV engineering approaches to enhance EV cell/tissue targeting and cargo delivery are highly desirable [23,28,29]. In relation to these challenges the global EV research community has also highlighted the urgent need for potency assays that can attempt to decipher EV therapeutic modalities and enable the measurement of EV therapeutic efficacy [2,30]. These and other EV research challenges are desirable to overcome, so that proposed therapeutic EV mechanisms are validated and more clinically effective therapeutic EVs can be developed.

We envision that innovative solutions to these challenges may reside at the convergence of EV research with engineering biology [16,31,32]. Indeed, it is already the case that EV studies are increasingly adopting synthetic biology methods and technologies to engineer cells, EVs and their cargoes, including CRISPR/Cas, SpyCatcher, fusion-protein engineering, and artificial intelligence (AI) approaches (e.g., AlphaFold) [29,31,33–35]. However, despite these recent advancements, it is still the case that cell-based engineering is relatively slow and is encumbered by cell growth and burden dynamics that can extend workflows across several days, weeks or even months [36,37]. In contrast, cell-free synthetic biology encompasses a suite of *in vitro* technologies, flexible engineering approaches and reaction formats that are rapid, spanning only several hours in duration [38–40]. Essentially, cell-free systems make use of isolated cellular components (proteins, nucleic acids etc.) or biological processes outside of the cell, and can, for example, enable *in vitro* protein production from DNA templates [40–43]. To achieve this, cell-free gene expression systems typically make use of isolated cellular transcription and translation machinery (e.g., protein synthesis using recombinant elements [PURE]) [44–46], or whole-cell extracts [47–51]. Given the unique biochemistries and protein synthesis yields of different cell-types, various cell-extract based systems have been developed, each with their own distinctive capabilities [40,52–55]. Cell-free reactions can also be further customised through supplementation with reaction-boosting energy mixes, enzyme co-factors, chaperones and other application-specific components as required [56–59]. For example, the inclusion of lipid nanostructures (e.g., liposomes, nanodiscs and bicelles) can facilitate the correct folding and insertion of cell-free produced membrane proteins [60–63]. Lipid-integrated membrane proteins can then be used for structural biology [64,65], drug screening [59,66], or synthetic cell applications [67,68].

These and other types of cell-free synthetic biology workflows can generate insights comparable to those obtained using cell-based characterisation assays [38–40,48,69]. Indeed, interoperability between cell-free and cell-based systems has enabled cell-free synthetic biology to underpin key advances in molecular biology, metabolic engineering, biosensing, materials research, drug discovery, therapeutic biomanufacturing and several areas of EV research [25,32,38,40,60,70–74]. For example, a cell-free EV packaging assay was developed that helped uncover a role for Y-box protein 1 in the sorting of specific mircoRNAs into exosomes [25]. Similar cell-free approaches could also be developed to study other aspects of EV biogenesis and cargo loading [32]. Whilst in other studies, cell-free gene expression systems have been used to produce many different recombinant proteins, including those of relevance to EV biology [32,60,70,71,75]. Inspired by these disparate studies, we demonstrated that it is also feasible to produce membrane fusion proteins in cell-free reactions that can co-localise and integrate with exogenously added HEK293 cell EVs [32]. In the future, we envision that this type of cell-free EV engineering could become an important strategy to rapidly prototype and test many different *in vitro* engineered EV designs for foundational or translational research aims [32]. We imagine applying cell-free-based EV engineering to rapidly decorate cell-derived or synthetic EVs with libraries of vaccine, therapeutic, cell-targeting or other elements for testing in downstream potency assays [16,32,76]. Importantly, this would allow for more systematic and engineering biology-based approaches to EV research, by making it easier for interlaboratory collaboration and data sharing [31]. Furthermore, cell-free optimised designs might then be transferred into cell-based EV engineering workflows to facilitate final testing and manufacturing scale up.

Here we present progresses beyond our earlier study [32] to further advance cell-free EV engineering as a viable EV research tool. Specifically, we demonstrate that hollow fibre-based cell culture is a feasible and scalable way to generate significant amounts of EVs for cell-free EV engineering workflows. Notably, hollow fibre-based bioreactors are increasingly being utilised in industry to biomanufacture clinical grade EV therapeutics. Additionally, our data highlights the impact that different EV isolation methods can have on EV batch characteristics and on downstream cell-free EV engineering performance. We also demonstrate that different membrane protein scaffold designs impact cell-free EV engineering efficiency, and that these observed differences can be readily measured using nano flow cytometry. We anticipate that this present study will add to future progress and improvements in cell-free EV engineering approaches and clinical applications.

## Results and discussion

### Hollowfibre biomanufacturing of HEK293 extracellular vesicles

Hollowfibre cell culture systems can facilitate high-density cell culture and scalable EV biomanufacturing that is compliant with good manufacturing practice (GMP) [22,77,78]. Hollowfibre bioreactors consist of a hollowfibre cartridge, within which cells are cultured, aided by a cell media reservoir, a gas exchanger, and a pump to facilitate continuous circulating flow of cell media throughout the bioreactor (Figure 1A). Hollowfibre cartridges are typically compact in size and contain many small fibres (∼0.2-1 mm in diameter) that collectively provide a large surface area, upon which cultured cells can adhere. Hollowfibre cartridge fibres are also porous, containing small holes that enable fresh cell media to pump through the entire cartridge, thereby providing nutrients and oxygen to the cultured cells in a manner that mimics *in vivo* circulation. Usefully, fibre pore sizes can be configured to ensure that cells and their secreted products, including EVs, of a desired molecular weight cut-off remain highly concentrated within the cartridge. Once established cells can remain in continuous culture for several weeks or more, whilst simultaneously the EV-enriched cell conditioned media can be readily harvested for downstream processing.

**Figure 1.**
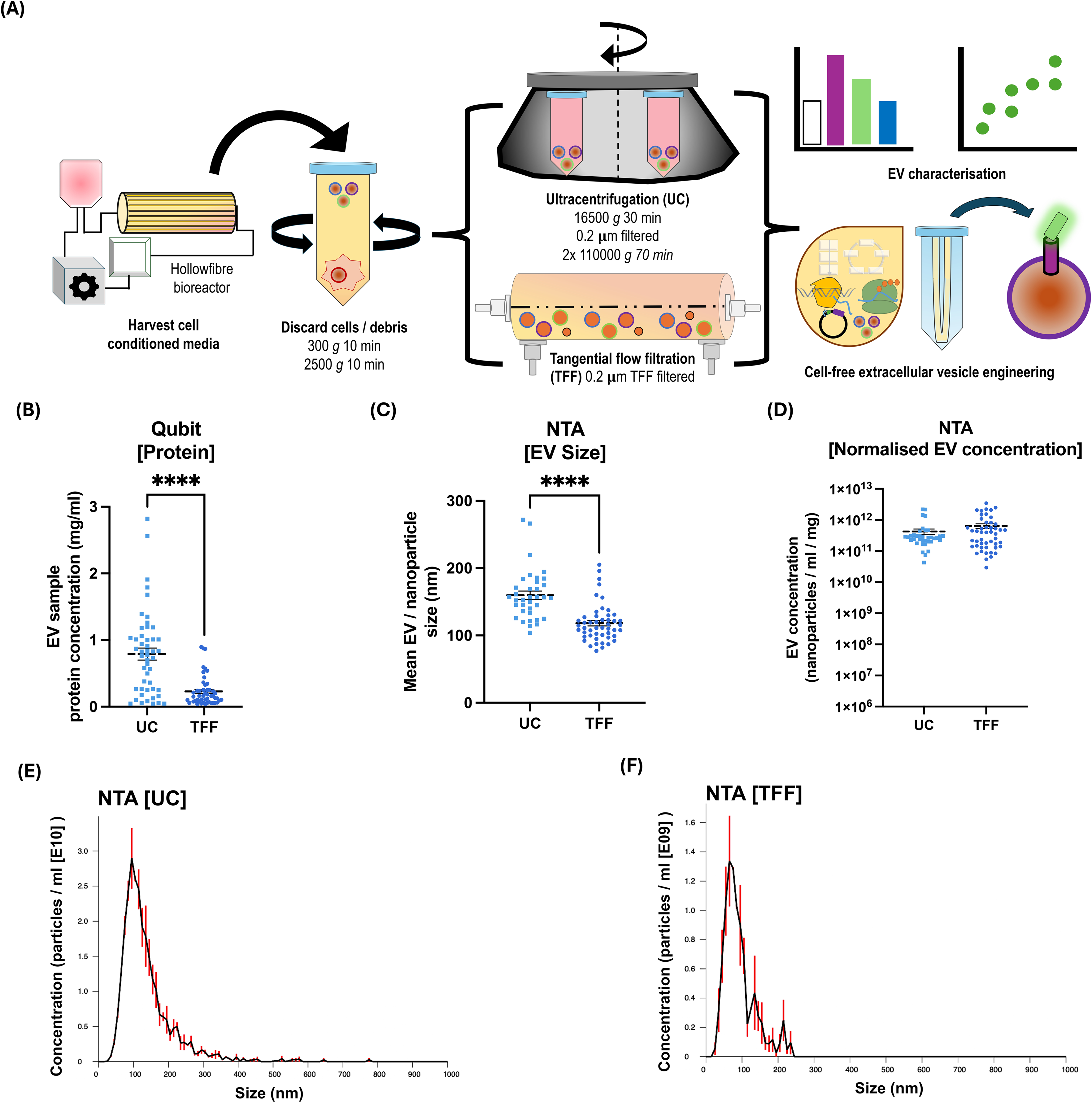
Hollowfibre biomanufacturing of HEK293 extracellular vesicles. **(A)** Workflow schematic depicting hollow-fibre HEK293 cell culture, extracellular vesicle (EV) isolation and cell-free EV engineering. Thirty-six independent batches of HEK293 cell conditioned media were harvested from the hollowfibre bioreactor and from these, HEK293 EVs were isolated using either ultracentrifugation (UC) or tangential flow filtration (TFF). EV batches were quantified for total protein concentration using **(B)** Qubit protein assay and nanoparticle tracking analysis (NTA) to quantify **(C)** EV size and **(D)** EV concentration, which was normalised according to batch total protein concentration. Representative NTA histograms for **(E)** UC and **(F)** TFF-isolated HEK293 EVs. Error bars denote standard error of the mean. Error bars denote standard error of the mean, *n* = 49 EV batches, student *t*-test **** *P*<0.0001.

To test the use of a hollowfibre bioreactor for EV production, we continuously cultured HEK293 cells for 26 days within a FiberCell Systems hollowfibre bioreactor equipped with a 20□kDa molecular weight cut-off cartridge and a 1 L media reservoir, that enabled cell culture densities of up to ∼10^9^ HEK293 cells. Similar to previous studies, cell conditioned batches (up to 20 ml each) were harvested from the hollowfibre cartridge and initially processed (centrifuged) to remove cells and large debris prior to EV isolation (Figure 1A) [8,22,32]. EVs were subsequently isolated from each cell conditioned media batch using ultracentrifugation (UC) and tangential flow filtration (TFF) approaches as shown in Figure 1A, (see materials and methods). Whilst HEK293 EVs were isolated using UC and TFF from the same cell conditioned media batches, there were differences. Notably, the average protein concentrations of UC-isolated EV samples (0.79 ±0.09 mg/ml) were higher than TFF-isolated EV samples (0.22 ±0.03 mg/ml) (Figure 1B), which are likely indicative of co-isolated contaminating proteins.

Post protein quantification, UC and TFF-isolated HEK293 EV samples were characterised using a nanoparticle tracking analysis (NTA) system to determine mean EV/nanoparticle sizes and concentrations (Figures 1C-F; Supplementary Figures 1 & 2). Mean EV/nanoparticle sizes of UC-isolated EVs were 159.8 ±6.23 nm, whilst TFF-isolated EV/nanoparticles were on average slightly smaller at 118.2 ±4.04 nm (Figure 1C), though these size ranges are both consistent with widely reported small EV sizes (< 200 nm) [2,30]. EV sizes were also measured using dynamic light scattering (DLS) and were broadly consistent with our NTA data. DLS mean EV/nanoparticle sizes for UC samples were 186 ±10.09 nm and for TFF samples were 130.2 ±16.66 nm (Supplementary Figure 3). For comparative purposes NTA-quantified EV/nanoparticle concentrations were normalised against sample protein concentration and sample volume differences (UC 200 μl; TFF 1000 μl; See Supplementary Figure 1 for pre-normalised data). Representative NTA-quantified EV size and concentrations for UC and TFF-isolated EVs are shown in Figures 1E and 1F respectively. Normalised mean EV/nanoparticle concentrations of UC-isolated HEK293 EVs were 4.27 ±0.86 x 10^11^ / ml and for TFF-isolated samples were 6.39 ±1.1 x 10^11^ / ml (Figure 1D). In addition to producing higher EV yields, we and other studies have shown that TFF protocols are simpler, quicker and more scalable than UC-based EV isolation protocols [79–82]. Nevertheless, both UC and TFF EV isolation methods generated sufficient HEK293 cell-derived EVs for further downstream characterisation and cell-free EV engineering experiments.

### Characterisation of HEK293 extracellular vesicles

In addition to NTA and DLS analyses, we further characterised several batches of UC and TFF-isolated HEK293 EVs using Minimal Information for Studies of Extracellular Vesicles (MISEV)- compliant methodologies [83]. Firstly, EV and cellular protein markers were assessed in four independent batches of UC and TFF-isolated HEK293s EVs, using an Exo-Check exosome antibody dot-blot array (Figures 2A & 2B; Supplementary Figure 4). UC and TFF-isolated EV batches were visibly positive for several EV markers including CD63, CD81 and TSG101 (Figures 2A & 2B; Supplementary Figure 4) [83], whilst all tested UC and TFF EV samples were strongly positive for the EV marker ALIX [83,84] (Supplementary Figure 4). Finally, we observed that GM130, a cis-Golgi matrix protein, was more consistently strongly positive in UC than TFF-isolated EV samples (Supplementary Figure 4), suggesting that these samples contain higher levels of cellular protein contaminants. However, this is not unexpected given previous studies also observed that UC methods invariably produce impure EV samples [79,85]. These data are also consistent with our previous observations in which UC samples were generally higher in overall protein concentration than TFF-isolated EV samples (Figure 1B), despite having lower normalised EV concentrations (Figure 1D). Nevertheless, these data confirm that both methods isolate EVs that display several common markers.

**Figure 2.**
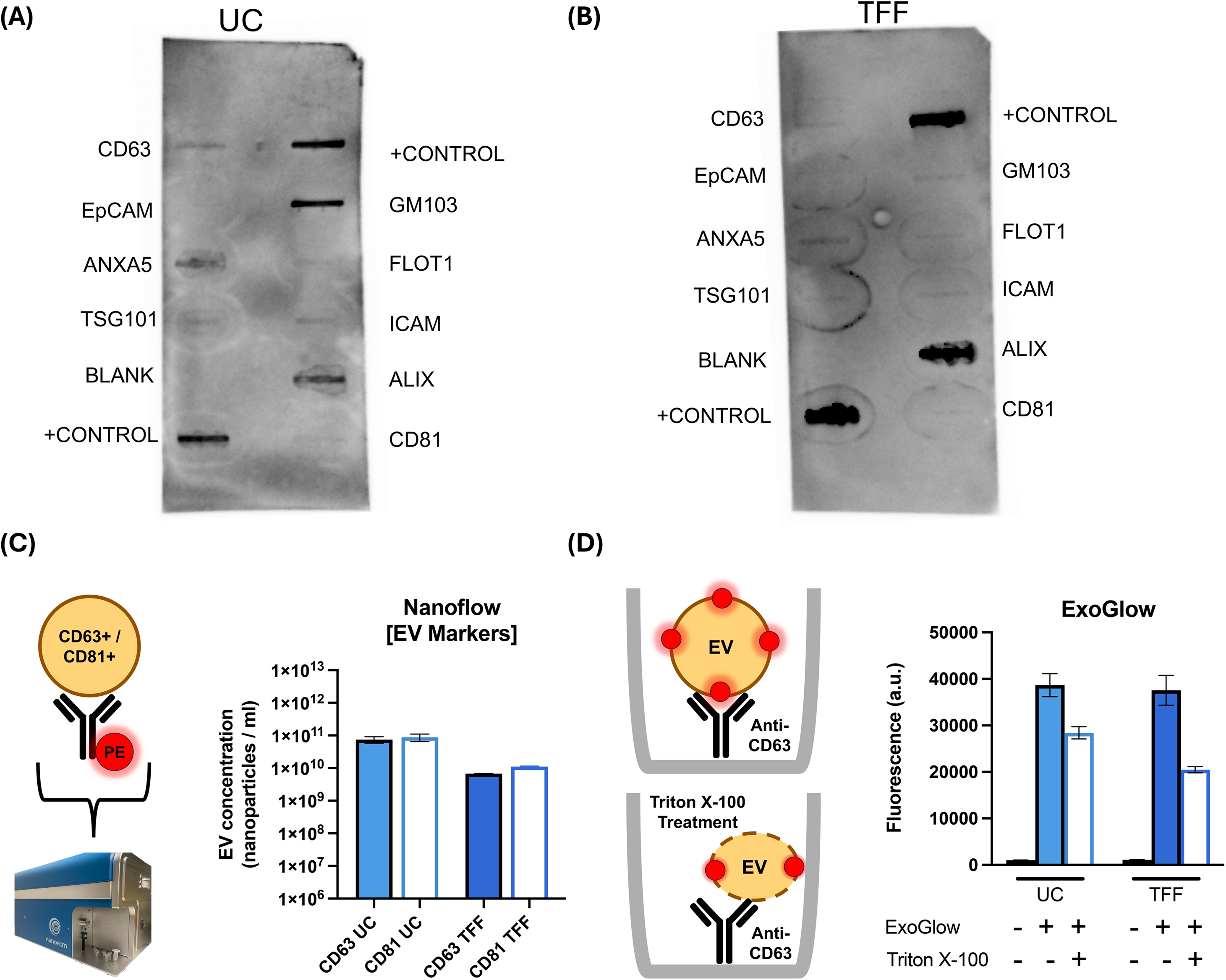
Characterisation of HEK293 extracellular vesicles. Representative Exo-Check exosome antibody arrays of **(A)** ultracentrifugation (UC) and **(B)** tangential flow filtration (TFF) isolated HEK293 extracellular vesicles. **(C)** Nano flow cytometry characterisation of CD63 and CD81 positive HEK293 EVs isolated using either UC or TFF, n=4 **(D)** ExoGlow EV membrane labelling of CD63 positive HEK293 EVs treated with 0 or 0.1% Triton X-100. Assay plates were analysed using a CLARIOstar plate reader (Ex:465-15/Em:635-20), n=4, Error bars denote standard error of the mean.

Representative EV samples were also imaged using super resolution microscopy, using a microscope equipped for direct stochastic optical reconstruction microscopy (dSTORM). Our images visually confirmed that UC and TFF-isolated HEK293 EVs were heterogeneously positive for the EV marker tetraspanins CD63, CD81 and CD9 (Supplementary Figure 5). CD63 and CD81 positive EVs were also detected using nano flow cytometry (Figure 2C; Supplementary Figures 6 & 7). Prior to use, the nano flow was calibrated using QC and size reference standard beads, to validate machine performance and to calibrate nanoparticle scatter measurements to EV/nanoparticle size (Supplementary Figure 6). We then stained UC and TFF-isolated HEK293 EVs with antibodies for CD63 or CD81 and then analysed using the calibrated nano flow cytometer (Figure 2C; Supplementary Figures 6 & 7). Mean EV/nanoparticle sizes of UC (69 ±0.76 nm) and TFF (78 ±1.3 nm) isolated EVs were smaller than our NTA and DLS measurements, though these data are consistent with nano flow cytometry measurements of HEK293 EVs in prior studies [20,32]. For the final EV characterisation assay, UC and TFF-isolated HEK293 EVs were immunocaptured using a CD63-antibody coated plate and stained with an ExoGlow membrane labelling dye to detect for intact EVs (Figure 2D). As expected, EV membrane integrity and concomitant ExoGlow dye fluorescence were reduced following Triton X-100 treatment (Figure 2D), thereby indicating that structurally intact CD63-positive EVs were present in the tested UC and TFF-isolated EV batches. Once characterised, several EV batches were ready for subsequent cell-free EV engineering experiments.

### Cell-free extracellular vesicle engineering

For cell-free EV engineering we constructed a panel of cell-free expression constructs cloned into pT7CFE1-Chis mammalian cell-free expression vector (STAR Methods, Supplementary Table 1). The choice of pT7CFE1-Chis was due to the plasmid also encoding a T7 RNA polymerase promoter (pT7) and an internal ribosomal entry site (IRES) from the encephalomyocarditis virus (EMCV), that can facilitate cell-free gene expression of cloned inserts in HeLa and other eukaryotic-based cell-free systems [32,49,50]. This new panel included the enhanced green fluorescent protein deGFP [86] and monomeric green lantern (mGL) [87], as reporter constructs deGFP (pRK66) and mGL (pRK67), respectively. Full-length CD63 (tetraspanin) and its truncated variant CD63ITM3 (CD63 I-shaped membrane-insertion topology transmembrane helix 3) were fused using a short flexible amino acid linker with mGL, to create the cell-free EV engineering constructs CD63-mGL (pRK68) and CD63ITM3-mGL (pRK69). CD63 and its truncated form CD63ITM3 were identified as suitable membrane proteins for cell-free EV engineering experiments, due to their demonstrated associations with exosomes in cell-based EV engineering studies [35]. Additionally, as we observed previously, cell-free produced EGFP-CD63 can also co-localise with HEK293 EVs [32]. It was therefore anticipated that these new CD63-based membrane fusion proteins would be suitable for cell-free EV engineering experiments.

We first sought to determine whether UC or TFF-isolated HEK293 EVs can impact HeLa cell-free reaction performance (Supplementary Figure 8). Briefly, 10 μl-scale cell-free reactions containing different HEK293 EV concentrations were setup in 384-well plates, and time course fluorescence measurements were used to assess cell-free deGFP production (Supplementary Figures 8 & 9). As expected, deGFP cell-free reactions produced ∼77 ±0.67 μg/ml of deGFP whilst control cell-free reactions containing neither deGFP DNA nor HEK293 EVs produced only low background levels of fluorescence (Supplementary Figure 8). Interestingly, the inclusion of UC or TFF-isolated HEK293 EVs modestly improved cell-free reaction performance, though cell-free reactions containing low EV concentrations produced more deGFP than reactions containing high EV concentrations (Supplementary Figure 8). These observed differences in cell-free deGFP production yields were most pronounced between cell-free reactions containing low (∼94 ±0.68 μg/ml) and high (∼83 ±1.75 μg/ml) concentrations of UC-isolated HEK293 EVs. Similarly to this, we previously observed that the inclusion of UC-isolated HEK293 EVs can negatively impact cell-free protein synthesis yields [32]. In contrast, TFF-isolated HEK293 EVs appear to have less pronounced impacts on cell-free reaction performance than UC-isolated HEK293 EVs (Supplementary Figure 8), which might be due to the relatively lower levels of protein contaminants in these TFF-isolated EV samples (Figures 1 & 2).

Given this, future cell-free EV engineering studies could explore whether improved TFF or other non-UC-based EV isolation methods are able to generate purer EV preparations, that have more consistent impacts on cell-free reaction performance. Indeed, EVs can contain chaperones, enzymes and other molecules that can impact cell-free metabolic processes and protein synthesis [32]. Nevertheless, in the present study both UC and TFF-isolated HEK293 EVs positively impacted cell-free reaction performance and were used for our cell-free EV engineering reactions.

Cell-free engineering of HEK293 EVs was carried out within semi-continuous dialysis mode cell-free reactions (Figure 3 & Supplementary Figure 10), using a protocol that we adapted from our previous study [32]. Essentially, reactions were setup for cell-free production of mGL, CD63-mGL or CD63ITM3-mGL proteins in the presence of UC or TFF-isolated HEK293 EVs, whilst control reactions included both UC and TFF-isolated HEK293 EVs but did not include a cell-free expression construct. The reactions were incubated at 30°C for 22 h, as described in Materials and Methods. Post-incubation, visual inspections (Figure 3A), and endpoint fluorescence measurements (Figure 3B) were indicative that mGL, CD63-mGL and CD63ITM3-mGL proteins were successfully produced in their respective cell-free reactions.

**Figure 3.**
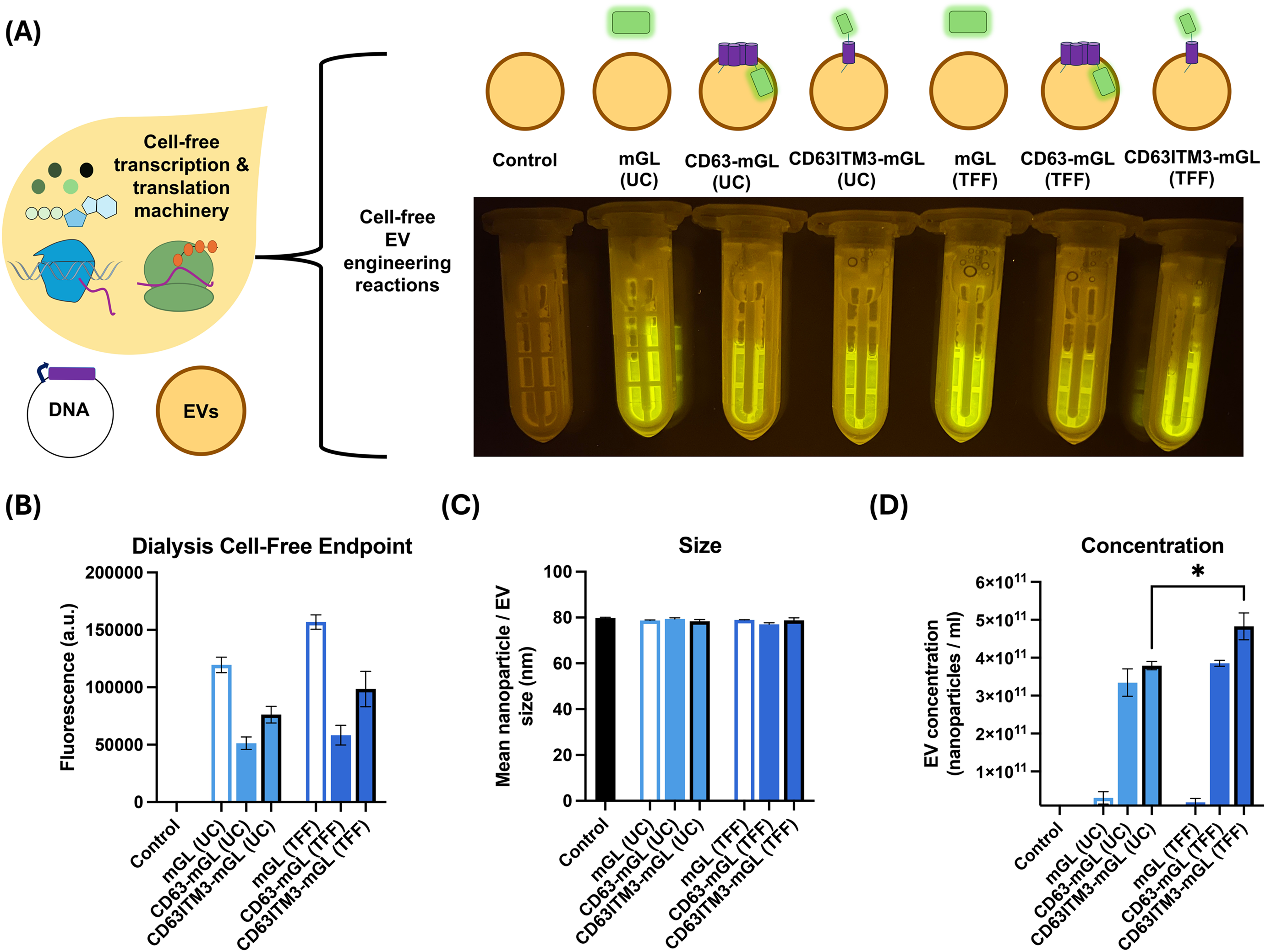
Cell-free engineering of HEK293 extracellular vesicles. **(A)** Schematic of ultracentrifugation (UC) and tangential flow filtration (TFF) isolated HEK293 EVs in control (no DNA), monomeric green lantern (mGL), CD63-mGL membrane fusion protein or truncated CD63ITM3-mGL membrane fusion protein producing cell-free EV engineering reactions. Representative 60 μL-scale ‘semi-continuous dialysis mode’ cell-free transcription-translation reactions were placed on a BluPAD LED transilluminator (Bio-Helix) and imaged as shown. **(B)** Endpoint (22 h) fluorescence measurements of control (no DNA), mGL, CD63-mGL or CD63ITM3-mGL protein production in 60 μL-scale ‘semi-continuous dialysis mode’ cell-free EV engineering reactions, containing UC or TFF-isolated HEK293 EVs. Endpoint samples were analysed using a CLARIOstar plate reader (Ex:483-14/Em:530-30). Nanoflow cytometry analysis of **(C)** mean nanoparticle/EV size and **(D)** concentration of green-fluorescent positive cell-free engineered HEK293 nanoparticles/EVs within the indicated ‘semi-continuous dialysis mode’ cell-free gene expression reactions. Error bars denote standard error of the mean, *n* = 3 independent cell-free reactions, student *t*-test \**P*□<□0.05.

These cell-free EV engineering reactions were subsequently processed and prepared for nano flow cytometry analyses, which confirmed the presence of nanoparticles/EVs in all samples (Figure 3C). Gated nanoflow cytometry events in all CD63-mGL and CD63ITM3-mGL cell-free EV engineering reactions indicate relatively high concentrations of strongly green fluorescent (FITC-H) nanoparticles/EVs (Figure 3D & Supplementary Figure 10). Interestingly, higher concentrations of green fluorescent positive nanoparticles/EVs were present in cell-free reactions containing TFF than UC-isolated HEK293 EVs (Figure 3D). Indeed, the highest concentrations of green fluorescent nanoparticles/EVs were found in CD63ITM3-mGL TFF cell-free reactions which were ∼4.83 ±0.35 ×10^11^ /ml (Figure 3D). The negative control reactions showed no fluorescence and only ∼0.4% of nanoparticles/EVs were green fluorescent in mGL cell-free EV engineering reactions (Figure 3D & Supplementary Figure 10). Consistent with these results, an anti-GFP-AF647 antibody displayed negligible staining of nanoparticles/EVs from control or mGL cell-free reactions (Supplementary Figure 10). Likewise, our observations of minimal mGL EV-surface aggregation are also consistent with our previous cell-free EV engineering study [32].

To further investigate cell-free EV engineering efficiencies, we also adapted our previous EV co-localisation and membrane protein topology assay [32] (Supplementary Figure 10). The purpose of the assay was to determine membrane protein topology given that correctly EV membrane integrated CD63-mGL would be green fluorescent (FITC-H) but would render any EV-internalised mGL inaccessible from GFP antibody staining (anti-GFP-AF647). Whilst EVs that have integrated CD63ITM3-mGL will be green fluorescent in a manner that might also be GFP antibody accessible (Supplementary Figure 10A). To this end, a quadrant-based nano flow cytometry gating strategy was developed to categorise recorded events according to their mGL (FITC-H) and GFP antibody fluorescence signals (anti-GFP-AF647) (Supplementary Figure 10B & C). These data were used to quantify cell-free EV engineering efficiencies of correctly integrated CD63-mGL (P4) and CD63ITM3-mGL (P1+P4) membrane fusion proteins with UC and TFF-isolated EVs (Supplementary Figure 10D). Interestingly, whilst we observed relatively high concentrations of strongly green fluorescent (FITC-H) nanoparticles/EVs in our cell-free EV engineering reactions (Figure 3D & Supplementary Figure 10), the percentage of events (nanoparticles/EVs) corresponding to correctly integrated membrane proteins was generally low (Supplementary Figure 10D). Though CD63ITM3-mGL co-localised and EV integrated more efficiently than CD63-mGL (Supplementary Figure 10D). This is perhaps unsurprising given the simpler single transmembrane architecture of CD63ITM3 compared with full length CD63. Nevertheless, UC and TFF-isolated EVs were successfully cell-free engineered with CD63-mGL and CD63ITM3-mGL. These data also offer an important insight in the development of additional membrane protein scaffolds that would likely further improve cell-free EV engineering capabilities and efficiencies.

### Concluding remarks

Globally EV research is growing rapidly to fully realise the potential of EVs as a powerful new class of vaccines, postbiotics and therapeutics. Although there are many foundational knowledge gaps in our understanding of EVs, for example biogenesis origins, and biological roles, translational EV research is accelerating to tackle key challenges. These include EV engineering, manufacturing and the need for EV potency assays that are currently hindering EV translational applications.

EV research is highly multidisciplinary and has already begun adopting engineering biology approaches including nascent examples of cell-free synthetic biology technologies. Cell-free synthetic biology encompasses several different *in vitro* technologies and in recent years has evolved into highly powerful prototyping platforms that can accelerate biotechnological development. Extrapolating these trends, we envision that it will likely become increasingly feasible to *in vitro* prototype, characterise, and improve the functionality (potency) of cell-free engineered EVs. In doing so, cell-free EV engineering could help accelerate future EV discoveries and the development of EV translational applications.

Here, we sought to further advance cell-free EV engineering by highlighting the importance of biomanufacturing and isolating EVs for cell-free EV engineering, using scalable GMP-compliant approaches. Prior to their use, our data also suggests that isolated EVs should be carefully characterised using MISEV-complaint EV characterisation methodologies, to understand batch heterogeneity and possible impacts on cell-free reaction performance. Furthermore, the engineering of additional membrane protein scaffolds would likely improve cell-free EV engineering capabilities and efficiencies. Finally, we also demonstrated that nano flow cytometry is a powerful tool for assessing cell-free EV engineering efficiency.

### Outstanding questions

- Are EVs from different cell types or disease states easier to cell-free engineer than others?
- Can insights on how different EV co-isolation contaminants impact cell-free reactions lead to the discovery of new accessory proteins that can further boost cell-free protein synthesis yields?
- Can cell-free EV engineering be used to generate sufficient EVs for downstream potency assays?
- Can insights from cell-free EV engineering be applied to adjacent fields, such as the development and engineering of synthetic cells?

## STAR Methods

### Key resources table

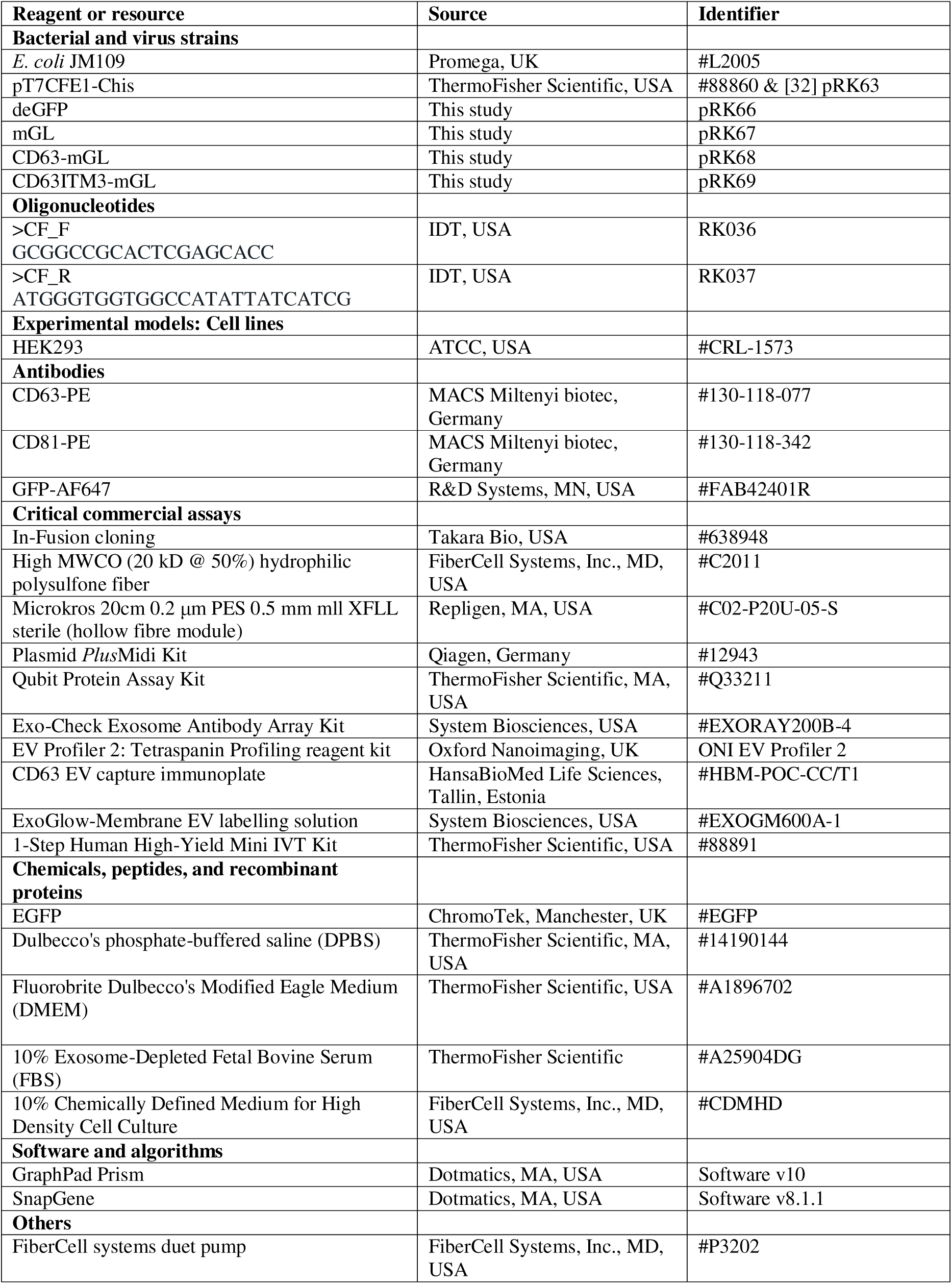
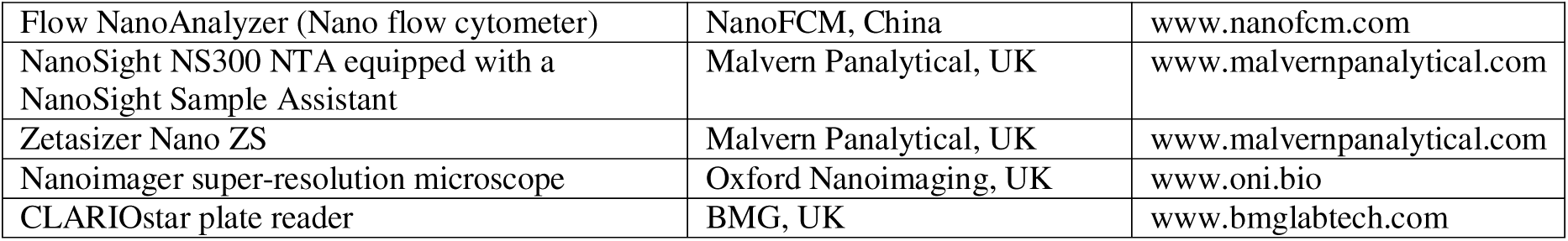

### Method Details

#### Bacterial strains and growth conditions

Plasmid constructs and strains used in this study are listed in the key resources table. *Escherichia coli* JM109 (#L2005, Promega, UK) was used for both cloning and generation of cell-free gene expression plasmids. For plasmid recovery *E. coli* strains were grown in Luria-Bertani (LB) medium supplemented with 100□μg/ml Ampicillin (w/v final concentration) and cultured at 37□°C with shaking (200□rpm).

#### Strain and plasmid construction

pT7CFE1-Chis (pRK63, #88860, ThermoFisher Scientific, USA) was commercially sourced from the 1-Step Human High-Yield Mini IVT Kit (#88891; ThermoFisher Scientific) and was transformed into competent *E. coli* JM109 (Promega) to create strain pRK63. For deGFP (pRK66), mGL (pRK67), CD63-mGL (pRK68) and CD63ITM3-mGL (pRK69) cell-free expression plasmids the inserts encoding the relevant proteins were designed using SnapGene cloning software (v8.1.1, Dotmatics, MA, USA) and then ordered as dsDNA gene synthesis fragments (gBlocks, IDT, USA). The plasmid backbone (pT7CFE1-Chis) was amplified and linearised using primer pair RK036 (CF_F) and RK037 (CF_R). PCR products were processed through agarose gel electrophoresis (1% w/v) and subsequently gel purified using the Monarch DNA gel extraction kit (#T1020, New England Biolabs, USA). Finally, In-Fusion cloning (Takara Bio, USA) of linearised pT7CFE1-Chis with the appropriate gBlocks (deGFP, mGL, CD63-mGL or CD63ITM3-mGL) followed by transformation into *E. coli* JM109 (Promega) was used to clone the indicated cell-free expression plasmids. Protein sequences are indicated in supplementary table 1. Constructs were verified using the whole-plasmid sequencing service provided by Full Circle Labs Ltd (London, UK).

To generate plasmids for cell-free experiments, glycerol stocks of the appropriate strains were used to inoculate 100 ml Luria-Bertani (LB) medium cultures supplemented with 100□μg/ml Ampicillin (final concentration), and these cultures were incubated overnight at 37□°C with shaking (200□rpm). Plasmid DNA was isolated using a Qiagen Plasmid *Plus*Midi Kit (#12943, Qiagen, Germany) and then ethanol precipitated prior to usage within the indicated cell-free reactions.

#### Hollow fibre cell culture

HEK293 cells (#CRL-1573, ATCC, USA) were cultured, as previously described [8,32], within a hollow fibre bioreactor (FiberCell Systems, Inc., MD, USA), which was configured with a 20□kDa molecular weight cut-off cartridge (#C2011, FiberCell Systems, Inc.) to enable high cell density culture (up to ∼10^9^ HEK293 cells) and the concentration of secreted HEK293 cell products (>20□kDa proteins and extracellular vesicles [EVs]) within the cartridge. Prior to cell seeding, the hollow fibre cartridge was sequentially primed, for 24 h each, with Dulbecco’s phosphate-buffered saline (DPBS, 1X; #14190144, ThermoFisher Scientific, MA, USA), then serum-free Fluorobrite Dulbecco’s Modified Eagle Medium (DMEM, #A1896702, ThermoFisher Scientific, USA) and finally Fluorobrite DMEM (#A1896702, ThermoFisher Scientific), supplemented with 10% Exosome-Depleted Fetal Bovine Serum (FBS, #A25904DG, ThermoFisher Scientific).

Once primed, the cartridge was seeded with 1×10^8^ adherent HEK293 cells that were cultured with Fluorobrite DMEM (#A1896702, ThermoFisher Scientific). Cell growth rate/viability was indirectly evaluated through daily monitoring of the glucose level of the cell culture medium (#GC001000, FiberCell Systems, Inc., MD, USA). The cell culture medium was changed when the glucose levels were below half the original level. Once glucose consumption levels significantly increased (50% glucose consumed within 24□h) the cell culture media was changed to Fluorobrite DMEM (#A1896702, ThermoFisher Scientific), supplemented with 10% Chemically Defined Medium for High Density Cell Culture (#CDMHD, FiberCell Systems). Up to two conditioned media samples (∼20□ml volume) were harvested from the hollow fibre cartridge each day.

#### Isolation of HEK293 extracellular vesicles

HEK293 cells were cultured in a hollow fibre bioreactor as described above and the harvested cell-conditioned media batches were split into two, such that one 5 ml aliquot was used to isolate extracellular vesicles using a previously described ultracentrifugation (UC) protocol [32,88] and the other 5 ml aliquot was used to isolate extracellular vesicles using tangential flow filtration (TFF). Briefly, 5 ml UC samples were first centrifuged at 300*g* for 10 minutes. The resultant supernatant was collected then centrifuged at 2,500*g* for 10 minutes. The supernatant was again collected and then further centrifuged at 16,500*g* for 30 minutes at 4°C with a Thermo Scientific Sorvall wX+ Ultra Series ultracentrifuge and T-647.5 fixed angle rotor. The supernatants were subsequently filtered using a 0.2 μm filter and then ultracentrifuged at 110000*g* for 70 minutes at 4°C. The resultant pellet was washed with 0.05 μm filtered DPBS (1X; #14190144, ThermoFisher Scientific), re-suspended in 0.05 μm filtered DPBS and then ultracentrifuged at 110,000*g* for 70 minutes at 4°C. Post-centrifugation, the supernatant was discarded, and the pellet containing extracellular vesicles was re-suspended into 200 μl of 0.05 μm filtered DPBS. In contrast, the 5 ml tangential flow filtration samples were first centrifuged at 300*g* for 10 minutes, then the supernatants were further centrifuged at 2,500*g* for 10 minutes to help remove cells and large debris. Prior to sample TFF-processing the sterile 0.2 μm tangential flow filtration modules (MicroKros #C02-P20U-05-S, Repligen, MA, USA) were sequentially pre-wetted prior to use with ethanol (20 % v/v) and DPBS (1X; #14190144; 0.05 μm filtered). Finally, centrifugation and TFF-processed cell media samples were normalised to 1 ml sample volumes.

### Extracellular vesicle characterisation

#### Qubit protein assay

The protein concentrations of ultracentrifugation (UC) and tangential flow filtration (TFF) isolated EV samples were determined, according to the manufactures’ guidance, using a Qubit 3 Fluorometer (Thermo Fisher Scientific, MA, USA) and a Qubit Protein Assay Kit (#Q33211, Thermo Fisher Scientific, MA, USA).

#### Nanoparticle tracking analysis (NTA)

UC-isolated HEK293 cell EVs were diluted (1:2000) and TFF-isolated HEK293 cell EVs were diluted (1:500) within particle-free DPBS (0.2 μm filtered; 1X; #14190144, Thermo Fisher Scientific), gently pipetted for 10□s and then aliquoted into a deep-well, 96-well plate. Samples from the 96-well plate were injected into a NanoSight NS300 NTA instrument equipped with a NanoSight Sample Assistant (Malvern Instruments, UK) autosampler system. The NTA images were recorded and analysed to obtain the concentration and distribution of the sample particles. Additional software version and measurement settings are shown in supplementary figure 2.

#### Dynamic light scattering (DLS)

UC and TFF-isolated HEK293 EVs were diluted (1:100) within particle-free DPBS (1X; 0.2 μm filtered; #14190144, Thermo Fisher Scientific), gently mixed by pipetting and transferred to a ZEN2112 quartz cuvette (Malvern Panalytical, Malvern, UK). Three technical replicates per sample were measured at 25□°C on a Malvern Zetasizer Nano ZS (Malvern Panalytical).

#### Exo-Check Exosome Antibody Array

Ultracentrifugation and tangential flow filtration isolated HEK293 EVs were lysed and processed, according to the manufacturer’s guidance, and analysed for the presence of a panel of EV markers using the Exo-Check Exosome Antibody Array Kit (#EXORAY200B-4, System Biosciences, USA). The Exo-Check array included 12 dot blot lines that incorporated antibodies specific for the EV markers CD63, CD81, ALIX, FLOT1, ICAM, EpCam, ANXA5 and TSG101, as well as a cellular contamination control (GM130 cis-Golgi marker) and several assay positive or negative [blank] controls. The dot blot array was developed using Immobilon Crescendo Western HRP substrate (#WBLUR0500, Millipore/Merck, Darmstadt, Germany) and imaged using a ChemiDoc imaging system (Bio-Rad Laboratories Inc., USA).

#### Super-resolution microscopy

Ultracentrifugation and tangential flow filtration isolated HEK293 extracellular vesicles were prepared and stained for super-resolution microscopy using an EV Profiler 2: Tetraspanin Profiling reagent kit for CD63, CD81 and CD9 tetraspanins, according to manufacturer’s recommendations (Oxford Nanoimaging [ONI], UK). EV samples were imaged using direct stochastic optical reconstruction microscopy (dSTORM), using a Nanoimager super-resolution microscope (Oxford Nanoimaging [ONI]) using laser channels (CD63-561, CD81-647 & CD9-488) and settings shown in Supplementary Figure 5. Resultant images were processed and analysed using the CODI online platform (Oxford Nanoimaging [ONI] https://alto.codi.bio/).

#### Nano flow cytometry analysis of EV markers

100 μl EV staining samples were setup including 3 μl of ultracentrifugation or 3 μl of tangential flow filtration isolated HEK293 extracellular vesicles, 0 μl or 5 μl of either CD63-PE (#130-118-077, MACS Miltenyi biotec, Germany) or CD81-PE (#130-118-342, MACS Miltenyi biotec, Germany) and 92 μl or 97 μl of 0.2 μm filtered DPBS (1X; #14190144, ThermoFisher Scientific). EV staining samples were pipette mixed and incubated at room temperature (21°C) for 30 minutes, with 350 rpm shaking (Thermo Mixer C, Eppendorf, Germany). Post staining, these 100 μl EV staining samples were processed using Vesi-SEC micro size exclusion chromatography columns (Vesiculab, Nottingham, UK) to remove unbound antibody. Briefly, Vesi-SEC micro columns were centrifuged at 1,000*g* for 1 minute to remove storage buffer, then post-centrifugation were placed into clean 1.5 ml tubes. 100 μl EV staining samples were loaded into the prepared Vesi-SEC micro columns and incubated for 1 minute. Post incubation, samples were centrifuged at 1,000*g* for 1 minute and the eluates (stained EV samples) were diluted 1:825 (UC) or 1:297 (TFF) within 0.2 μm filtered DPBS (1X; #14190144, ThermoFisher Scientific) ready for nano flow cytometry analysis (Flow NanoAnalyzer, NanoFCM, China). The Flow NanoAnalyzer was calibrated using QC Beads (250 nm SiNP Dual laser QC Beads NanoFCM #QS2503) and size reference standard beads (Silica nanospheres, NanoFCM #S16M-Exo 68-155 nm) were used to calibrate the size distribution of EVs, whilst 0.2 μm filtered DPBS (1X; #14190144, ThermoFisher Scientific) was used as a blank control. Standards and samples were recorded for 1 minute using excitation laser 488 nm, SSC (488/10 nm) and PE (580/40 nm) filters. Data analysis (including sample thresholding to remove background and detector noise signals) was carried out, according to the manufacturer’s recommendations and best practices [89], using Flow NanoAnalyzer software (v2.0).

#### ExoGlow Membrane EV labelling

Control and EV samples were setup for ExoGlow labelling with 0 μl or 5 μl of HEK293 extracellular vesicles (2.5×10^10^ / ml of either ultracentrifugation or tangential flow filtration isolated EVs as indicated) and 81 μl or 86 μl of either 0.2 μm filtered DPBS (1X; #14190144, ThermoFisher Scientific) or 0.2 μm filtered DPBS-T (DPBS with 0.1% [v/v] Triton X-100). These samples were incubated for 10 minutes at room temperature (21°C) and then added to appropriate wells in an CD63 EV capture immunoplate (#HBM-POC-CC/T1, HansaBioMed Life Sciences, Tallin, Estonia). Appropriate wells were stained with either 0 μl or 14 μl of ExoGlow-Membrane EV labelling solution (2 μl labelling dye and 12 μl reaction buffer; #EXOGM600A-1, System Biosciences), gently pipette mixed and incubated at room temperature (21°C) in the dark for 60 minutes with 100 rpm shaking (Thermo Mixer C, Eppendorf, Germany). Post incubation, wells were gently washed twice with 100 μl of 0.2 μm filtered DPBS (1X; #14190144, ThermoFisher Scientific) and then filled with 100 μl of 0.2 μm filtered DPBS (1X; #14190144, ThermoFisher Scientific). Plates were measured on a CLARIOstar plate reader (BMG, UK) with the following settings - excitation 465-15 nm, 548.8 nm dichroic filter and emission 635-20 nm with 1800 gain.

#### Cell-free time course reactions

Cell-free time course reactions (10 μl) were prepared using a 1-Step Human High-Yield Mini IVT Kit (#88891; ThermoFisher Scientific, USA) and consisted of the following components: 2 μl Reaction Mix, 1 μl Accessory Proteins, 5 μl HeLa cell lysate, 0 or 400 ng of plasmid DNA (deGFP), 0 μl, 0.3 μl [Low] or 0.6 μl [High] of HEK293 cell EVs (2.5×10^10^ / ml of either ultracentrifugation [UC] or tangential flow filtration [TFF] isolated HEK293 EVs as indicated) and nuclease free water to top up the reactions to 10 μl (final volume). These 10 μl cell-free time course reactions were pipetted into individual wells of 384-well plates (Griener bio-one, NC, USA) and measured using a CLARIOstar plate reader (BMG, UK) with the following settings: 30°C incubation temperature, excitation laser 483-14 nm, 502.5 dichroic and emission filter 530–30 nm with 1000 gain. Plates were sealed and incubated within the plate reader at 30°C. Sample plates were shaken (500 rpm) for 5 seconds prior to each 10 min reading cycle during the 10 h time course. Endpoint measurements were taken at 200 minutes and were used, along with a GFP calibration curve, to calculate GFP yields in the respective cell-free time course reactions. The GFP calibration curve was adapted from previous studies [8,48].

Briefly, EGFP (#EGFP; ChromoTek, Manchester, UK) was serially diluted 0-1 mg/ml, aliquoted as 10 ul volumes in technical quadruplicate into appropriate wells of a 384-well plate (#781096, Greiner Bio-One, Kremsmünster, Austria) and measured using a CLARIOstar plate reader (excitation laser 483-14 nm, 502.5 nm dichroic and emission filter 530–30 nm with 1000 gain). These data were used to plot the GFP calibration curve (Supplementary Figure 9) using GraphPad Prism v10 software

#### Dialysis-mode cell-free EV engineering reactions

Cell-free reaction master mixes (60 μl) were prepared using a 1-Step Human High-Yield Mini IVT Kit (#88891; ThermoFisher Scientific, USA) and consisted of the following components: 12 μl Reaction Mix, 6 μl Accessory Proteins, 30 μl HeLa cell lysate, 2.4 μg of plasmid DNA (mGL, CD63-mGL or CD63ITM3-mGL), 3.6 μl of HEK293 EVs (2.5×10^10^ / ml of either control [1.8 μl UC + 1.8 μl TFF], ultracentrifugation [UC] or tangential flow filtration [TFF] isolated HEK293 EVs as indicated and nuclease free water to top up the master mixes to 60 μl. These 60 μl master mixes were aliquoted into a micro-dialysis cassette (10kDa MWCO, ThermoFisher Scientific/Pierce #88260) that was inserted into a 2 ml microtube pre-filled with 1.4 ml of Dialysis Buffer (from 1-Step Human High-Yield Mini IVT Kit #88891). Cell-free dialysis reactions were incubated in an Eppendorf Thermo Mixer C at 30°C with 950 rpm shaking for 22h. Post-incubation, these dialysis-mode cell-free EV engineering reactions were pipetted out from the dialysis cassettes and into clean 1.5 ml microtubes. 10 μl of these master mixes were subsequently aliquoted into individual wells of 384-well plates (Griener bio-one, NC, USA) and endpoint mGL fluorescence was measured using a CLARIOstar plate reader (BMG, UK) with the following settings: preheated to 30°C, excitation 483-14 nm, 502.5 dichroic and emission 530–30 nm with 1500 gain. The remaining samples were analysed using nano flow cytometry as described below.

#### Nano flow cytometry analysis of cell-free engineered EVs

Control and cell-free engineered HEK293 extracellular vesicles were antibody stained *in situ* (i.e. within their respective cell-free reactions) in staining reactions including 10 μl of the indicated cell-free EV engineering reaction (Control, mGL, CD63-mGL or CD63ITM3-mGL with UC and/or TFF-isolated HEK293 cell EVs), 90 μl of 0.2 μm filtered DPBS (1X; #14190144, ThermoFisher Scientific) and 0.5 μl GFP-AF647 conjugated antibody (0.1 μg; #FAB42401R, R&D Systems, MN, USA), totalling 100.5 μl (final volume). These EV staining samples were pipette mixed and incubated at room temperature (21°C) for 30 minutes with 350 rpm shaking (Thermo Mixer C, Eppendorf, Germany). Post-incubation, these EV staining samples were processed using Vesi-SEC micro size exclusion chromatography columns (Vesiculab, Nottingham, UK) to remove unbound antibody and to enrich for EVs. Briefly, Vesi-SEC micro columns were centrifuged at 1,000*g* for 1 minute to remove storage buffer, then the 100.5 μl antibody-stained cell-free engineered EV samples were loaded into the prepared Vesi-SEC micro columns, incubated for 1 minute and centrifuged at 1,000*g* for 1 minute to elute stained and non-antibody stained cell-free engineered EVs. Eluted EVs were diluted 1:16:000 within 0.2 μm filtered DPBS (1X; #14190144, ThermoFisher Scientific) ready for nano flow cytometry analysis (Flow NanoAnalyzer, NanoFCM, China). The Flow NanoAnalyzer was again calibrated using QC Beads (250 nm SiNP Dual laser QC Beads NanoFCM #QS2503) and size reference standard beads (Silica nanospheres, NanoFCM #S16M-Exo 68-155 nm), whilst 0.2 μm filtered DPBS (1X; #14190144, ThermoFisher Scientific) was used as a blank control. Standards and samples were recorded for 1 minute using 488 nm and 638 nm excitation lasers and several filters SSC (488/10 nm), FITC (525/40 nm) and AF647 (670/30 nm). Data analysis, including sample thresholding to remove background and detector noise signals was carried out, according to the manufacturer’s recommendations and best practices [89] using Flow NanoAnalyzer software (v2.0). Where indicated a quadrant gating strategy was used to identify cell-free engineered EVs (mGL +ve) and GFP antibody stained EVs (AF647 +ve).

## Resource availability

Data is included within the manuscript and supplementary files.

## Author contributions

RK: Conceptualization; Data curation; Formal analysis; Funding acquisition; Investigation; Methodology; Project administration; Resources; Supervision; Validation; Visualization; Writing—original draft; Writing—review & editing.

AJW: Conceptualization; Data curation; Formal analysis; Funding acquisition; Investigation; Methodology; Writing - review & editing.

AH: Data curation; Formal analysis; Investigation; Methodology; Writing—review & editing. PF: Conceptualization; Data curation; Formal analysis; Funding acquisition; Investigation;

Methodology; Project administration; Resources; Supervision; Validation; Visualization; Writing—original draft; Writing—review & editing.

## Acknowledgements

We thank colleagues in the Section of Structural and Synthetic Biology within the Department of Infectious Disease at Imperial College London for their advice and helpful comments. We also thank Pip Timmins and Franky Djutanta from ONI UK for their expertise and assistance with demoing of an ONI Nanoimager super resolution microscope. Our EV work was supported by grants from the Biotechnology and Biological Sciences Research Council (BBSRC) [BB/T017147/1], [BB/W012987/1], [UKRI2410], the Wellcome Trust [221546/Z/20/Z], the Engineering and Physical Sciences Research Council (EPSRC) and SynbiCITE proof of concept award [EP/Z533142/1], and we also thank the Future Biomanufacturing Research Hub (FBRH) at The University of Manchester and UKRI for a Future BRH Award [EP/S01778X/1].

## Supplemental information

Supplemental information to this article can be found online at [journal link]

## References

1. Couch, Y. et al. (2021) A brief history of nearly EVLerything – The rise and rise of extracellular vesicles. J Extracell Vesicles 10

2. Welsh, J.A. et al. (2024) Minimal information for studies of extracellular vesicles (MISEV2023): From basic to advanced approaches. J Extracell Vesicles 13

3. Cheng, K. and Kalluri, R. (2023) Guidelines for clinical translation and commercialization of extracellular vesicles and exosomes based therapeutics. Extracellular Vesicle 2, 100029

4. Mizenko, R.R. et al. (2024) A critical systematic review of extracellular vesicle clinical trials. J Extracell Vesicles 13

5. Kumar, M.A. et al. (2024) Extracellular vesicles as tools and targets in therapy for diseases. Signal Transduct Target Ther 9, 27

6. Dixson, A.C. et al. (2023) Context-specific regulation of extracellular vesicle biogenesis and cargo selection. Nat Rev Mol Cell Biol 24, 454–476

7. Holcar, M. et al. (2025) Comprehensive Phenotyping of Extracellular Vesicles in Plasma of Healthy Humans – Insights Into Cellular Origin and Biological Variation. J Extracell Vesicles 14

8. Kelwick, R.J.R.R. et al. (2021) AL-PHA beads: Bioplastic-based protease biosensors for global health applications. Materials Today 47, 25–37

9. Yim, K.H.W. et al. (2023) Assessing Extracellular Vesicles in Human Biofluids Using FlowLBased Analyzers. Adv Healthc Mater 12

10. Tran, H.L. et al. (2025) Extracellular Vesicles for Clinical Diagnostics: From Bulk Measurements to Single-Vesicle Analysis. ACS Nano 19, 28021–28109

11. Lener, T. et al. (2015) Applying extracellular vesicles based therapeutics in clinical trials – an ISEV position paper. J Extracell Vesicles 4, 30087

12. Colao, I.L. et al. (2018) Manufacturing Exosomes: A Promising Therapeutic Platform. Trends Mol Med 24, 242–256

13. Nieland, L. et al. (2023) Engineered EVs designed to target diseases of the CNS. Journal of Controlled Release 356, 493–506

14. Liu, Y.-J. and Wang, C. (2023) A review of the regulatory mechanisms of extracellular vesicles-mediated intercellular communication. Cell Communication and Signaling 21, 77

15. Erana-Perez, Z. et al. (2025) Differential protein and mRNA cargo loading into engineered large and small extracellular vesicles reveals differences in in vitro and in vivo assays. Journal of Controlled Release 379, 951–966

16. Kelwick, R.J.R. et al. (2023) Opportunities for engineering outer membrane vesicles using synthetic biology approaches. Extracell Vesicles Circ Nucl Acids 4, 255–61

17. Wang, Y. et al. (2025) Application of bacterial extracellular vesicles in gastrointestinal diseases. Trends Biotechnol DOI: 10.1016/j.tibtech.2025.05.022

18. Garling, A. et al. (2025) Outer Membrane Vesicles as a Versatile Platform for Vaccine Development: Engineering Strategies, Applications and Challenges. J Extracell Vesicles 14

19. Rezaie, J. et al. (2022) A review on exosomes application in clinical trials: perspective, questions, and challenges. Cell Communication and Signaling 20, 145

20. Chen, Z.-Q. et al. (2025) A comprehensive evaluation of stability and safety for HEK293F-derived extracellular vesicles as promising drug delivery vehicles. Journal of Controlled Release 382, 113673

21. Giebel, B. and Lim, S.K. (2025) Overcoming challenges in MSC-sEV therapeutics: insights and advances after a decade of research. Cytotherapy 27, 843–848

22. Gobin, J. et al. (2021) Hollow-fiber bioreactor production of extracellular vesicles from human bone marrow mesenchymal stromal cells yields nanovesicles that mirrors the immuno-modulatory antigenic signature of the producer cell. Stem Cell Res Ther 12, 127

23. Kim, H.I. et al. (2024) Recent advances in extracellular vesicles for therapeutic cargo delivery. Exp Mol Med 56, 836–849

24. Carney, R.P. et al. (2025) Harnessing extracellular vesicle heterogeneity for diagnostic and therapeutic applications. Nat Nanotechnol 20, 14–25

25. Shurtleff, M.J. et al. (2016) Y-box protein 1 is required to sort microRNAs into exosomes in cells and in a cell-free reaction. Elife 5, 1–23

26. Driedonks, T. et al. (2022) Pharmacokinetics and biodistribution of extracellular vesicles administered intravenously and intranasally to *Macaca nemestrina*. Journal of Extracellular Biology 1

27. Rosenkrans, Z.T. et al. (2024) Investigating the In Vivo Biodistribution of Extracellular Vesicles Isolated from Various Human Cell Sources Using Positron Emission Tomography. Mol Pharm 21, 4324–4335

28. Zhao, S., et al. (2024) Targeted delivery of extracellular vesicles: the mechanisms, techniques and therapeutic applications. Molecular Biomedicine 5, 60

29. Greenberg, Z.F. et al. (2023) Towards artificial intelligence-enabled extracellular vesicle precision drug delivery. Adv Drug Deliv Rev 199, 114974

30. Théry, C. et al. (2018) Minimal information for studies of extracellular vesicles 2018 (MISEV2018): a position statement of the International Society for Extracellular Vesicles and update of the MISEV2014 guidelines. J Extracell Vesicles 7, 1535750

31. Kelwick, R.J.R., et al. (2024) Accelerating extracellular vesicle research and biotechnological applications using synthetic biology approaches. Extracellular Vesicle 4, 100050

32. Kelwick, R.J.R. et al. (2023) Opportunities to accelerate extracellular vesicle research with cellLfree synthetic biology. Journal of Extracellular Biology 2

33. Zanella, I. et al. (2021) ProteomeLminimized outer membrane vesicles from *Escherichia coli* as a generalized vaccine platform. J Extracell Vesicles 10

34. Alves, N.J. et al. (2015) Bacterial Nanobioreactors–Directing Enzyme Packaging into Bacterial Outer Membrane Vesicles. ACS Appl Mater Interfaces 7, 24963–24972

35. Curley, N. et al. (2020) Sequential deletion of CD63 identifies topologically distinct scaffolds for surface engineering of exosomes in living human cells. Nanoscale 12, 12014–12026

36. Grob, A. et al. (2025) Design of an intracellular aptamer-based fluorescent biosensor to track burden in Escherichia coli. Trends Biotechnol DOI: 10.1016/j.tibtech.2025.06.007

37. Di Blasi, R., et al. (2023) Resource-aware construct design in mammalian cells. Nat Commun 14, 3576

38. Garenne, D. et al. (2021) Cell-free gene expression. Nature Reviews Methods Primers 1, 49

39. Chappell, J. et al. (2013) Validation of an entirely in vitro approach for rapid prototyping of DNA regulatory elements for synthetic biology. Nucleic Acids Res 41, 1–11

40. Hunt, A.C. et al. (2025) Cell-Free Gene Expression: Methods and Applications. Chem Rev 125, 91–149

41. Sun, Z.Z. et al. (2013) Protocols for implementing an Escherichia coli based TX-TL cell-free expression system for synthetic biology. J Vis Exp DOI: 10.3791/50762

42. Garenne, D. et al. (2021) The all-E. coliTXTL toolbox 3.0: new capabilities of a cell-free synthetic biology platform. Synth Biol 6

43. Kapasiawala, M. and Murray, R.M. (2024) Metabolic Perturbations to an *Escherichia coli*-based Cell-Free System Reveal a Trade-off between Transcription and Translation. ACS Synth Biol 13, 3976–3990

44. Shimizu, Y. et al. (2001) Cell-free translation reconstituted with purified components. Nat Biotechnol 19, 751–755

45. Lavickova, B. and Maerkl, S.J. (2019) A Simple, Robust, and Low-Cost Method To Produce the PURE Cell-Free System. ACS Synth Biol 8, 455–462

46. Zhang, Y. et al. (2025) Optimizing Protein Production in the One-Pot PURE System: Insights into Reaction Composition and Expression Efficiency. ACS Synth Biol 14, 1496–1508

47. Gagoski, D. et al. (2015) Benchmarking of four cell-free protein expression systems. Biotechnol Bioeng 9999, n/a-n/a

48. Kelwick, R. et al. (2016) Development of a Bacillus subtilis cell-free transcription-translation system for prototyping regulatory elements. Metab Eng 38, 370–381

49. Kopniczky, M.B. et al. (2020) Cell-Free Protein Synthesis as a Prototyping Platform for Mammalian Synthetic Biology. ACS Synth Biol 9, 144–156

50. Heide, C. et al. (2021) Design, Development and Optimization of a Functional Mammalian Cell-Free Protein Synthesis Platform. Front Bioeng Biotechnol 8

51. Wiegand, D.J. et al. (2018) Establishing a Cell-Free *Vibrio natriegens* Expression System. ACS Synth Biol 7, 2475–2479

52. Ogawa, A. et al. (2016) Rational optimization of amber suppressor tRNAs toward efficient incorporation of a non-natural amino acid into protein in a eukaryotic wheat germ extract. Org Biomol Chem 14, 2671–8

53. Moore, S.J. et al. (2018) Rapid acquisition and model-based analysis of cell-free transcription–translation reactions from nonmodel bacteria. Proceedings of the National Academy of Sciences 115

54. Sakai, A. et al. (2023) Cell-Free Expression System Derived from a Near-Minimal Synthetic Bacterium. ACS Synth Biol 12, 1616–1623

55. Meyerowitz, J.T. et al. (2024) Development of Cell-Free Transcription–Translation Systems in Three Soil Pseudomonads. ACS Synth Biol 13, 530–537

56. Borkowski, O. et al. (2020) Large scale active-learning-guided exploration for in vitro protein production optimization. Nat Commun 11, 1872

57. Cai, Q. et al. (2015) A simplified and robust protocol for immunoglobulin expression in E scherichia coli cell-free protein synthesis systems. Biotechnol Prog 31, 823–831

58. Olsen, M.L. et al. (2025) Design-driven optimization of low-cost reagent formulations for reproducible and high-yielding cell-free gene expression. DOI: 10.1101/2025.08.01.668204

59. Caschera, F. (2025) Cell-free protein synthesis platforms for accelerating drug discovery. Biotechnology Notes 6, 126–132

60. Kruyer, N.S. et al. (2021) Membrane Augmented Cell-Free Systems: A New Frontier in Biotechnology. ACS Synth Biol 10, 670–681

61. Shelby, M.L. et al. (2019) Cell-Free Co-Translational Approaches for Producing Mammalian Receptors: Expanding the Cell-Free Expression Toolbox Using Nanolipoproteins. Front Pharmacol 10

62. Jiang, S. et al. (2024) A cell-free system for functional studies of small membrane proteins. Journal of Biological Chemistry 300, 107850

63. Matsumoto, R. et al. (2025) Lipid Modification and Membrane Localization of Proteins in Cell-Free System. ACS Synth Biol 14, 2729–2738

64. Terada, T., et al. (2014) Cell-Free Expression of Protein Complexes for Structural Biologypp. 151–159

65. Köck, Z. et al. (2024) Cryo-EM structure of cell-free synthesized human histamine 2 receptor/Gs complex in nanodisc environment. Nat Commun 15, 1831

66. Zhu, J. et al. (2025) AI-driven high-throughput droplet screening of cell-free gene expression. Nat Commun 16, 2720

67. Giaveri, S. et al. (2025) Building a Synthetic Cell Together. Nat Commun 16, 7488

68. Cao, M. et al. (2024) Cell-free Protein Synthesis System for Building Synthetic Cells. Journal of Visualized Experiments DOI: 10.3791/66626

69. Kelwick, R. et al. (2018) Cell-free prototyping strategies for enhancing the sustainable production of polyhydroxyalkanoates bioplastics. Synth Biol 3

70. Shinoda, T. et al. (2016) Cell-free methods to produce structurally intact mammalian membrane proteins. Sci Rep 6, 30442

71. Umbach, S., et al. (2022) Cell-Free Expression of GPCRs into Nanomembranes for Functional and Structural Studiespp. 405–424

72. Kelwick, R.J.R. et al. (2020) Biological Materials: The Next Frontier for Cell-Free Synthetic Biology. Front Bioeng Biotechnol 8

73. Pardee, K. et al. (2014) Paper-based synthetic gene networks. Cell 159, 940–954

74. Pardee, K. et al. (2016) Portable, On-Demand Biomolecular Manufacturing. Cell 167, 248–259.e12

75. Sachse, R. et al. (2013) Synthesis of membrane proteins in eukaryotic cellLfree systems. Eng Life Sci 13, 39–48

76. Meyer, C. et al. (2025) Designer artificial environments for membrane protein synthesis. Nat Commun 16, 4363

77. Kink, J.A. et al. (2024) Large-scale bioreactor production of extracellular vesicles from mesenchymal stromal cells for treatment of acute radiation syndrome. Stem Cell Res Ther 15, 72

78. Garcia, S.G. et al. (2024) Hollow fiber bioreactor allows sustained production of immortalized mesenchymal stromal cell-derived extracellular vesicles. Extracell Vesicles Circ Nucl Acids 5, 201–20

79. Visan, K.S. et al. (2022) Comparative analysis of tangential flow filtration and ultracentrifugation, both combined with subsequent size exclusion chromatography, for the isolation of small extracellular vesicles. J Extracell Vesicles 11

80. Paolini, L. et al. (2022) LargeLscale production of extracellular vesicles: Report on the “massivEVs” ISEV workshop. Journal of Extracellular Biology 1

81. Onesti, R. et al. (2025) Continuous Tangential Flow Filtration to Isolate Extracellular Vesicles from Dairy By-Products. Ind Eng Chem Res 64, 15778–15789

82. Ng, K.S. et al. (2019) Bioprocess decision support tool for scalable manufacture of extracellular vesicles. Biotechnol Bioeng 116, 307–319

83. Welsh, J.A. et al. (2024) Minimal information for studies of extracellular vesicles (MISEV2023): From basic to advanced approaches. J Extracell Vesicles 13

84. Fordjour, F.K. et al. (2022) A shared, stochastic pathway mediates exosome protein budding along plasma and endosome membranes. Journal of Biological Chemistry 298, 102394

85. Bonner, S.E. et al. (2024) Scalable purification of extracellular vesicles with high yield and purity using multimodal flowthrough chromatography. Journal of Extracellular Biology 3

86. Shin, J. and Noireaux, V. (2010) Efficient cell-free expression with the endogenous E. Coli RNA polymerase and sigma factor 70. J Biol Eng 4, 8

87. Campbell, B.C. et al. (2020) mGreenLantern: a bright monomeric fluorescent protein with rapid expression and cell filling properties for neuronal imaging. Proceedings of the National Academy of Sciences 117, 30710–30721

88. Li, P. et al. (2017) Progress in Exosome Isolation Techniques. Theranostics 7, 789–804

89. Lees, R. et al. (2022) Single Extracellular Vesicle Transmembrane Protein Characterization by Nano-Flow Cytometry. Journal of Visualized Experiments DOI: 10.3791/64020

