## Supplementary for "Hollow-fibre biomanufacturing and cell-free engineering of HEK293 extracellular vesicles"

#Joint first authors

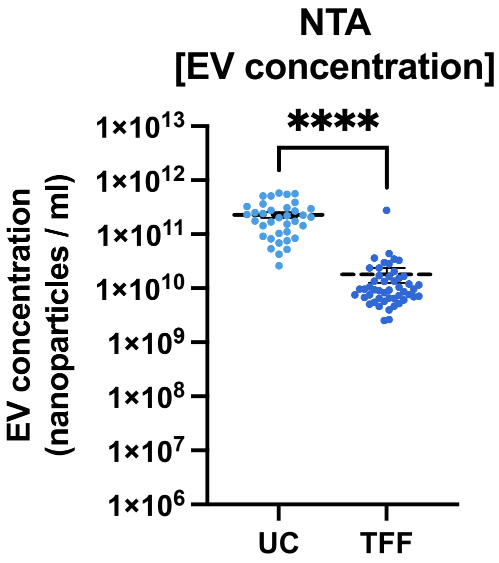


**Supplementary Figure 1. Nanoparticle tracking analysis of HEK293 extracellular vesicles (EVs) that were isolated using either ultracentrifugation (UC) or tangential flow filtration (TFF).** These data are prior to normalisation as shown in Figure 1D. Error bars denote standard error of the mean, $n$ = 49 EV batches, student $t$-test ****$P$<0.0001.

**(A)**

**Details**

| | |
|---|---|
| NTA Version: | NTA 3.4 - Sample Assistant Build 3.4.003-SA |
| Script Used: | SOP AutoSampler Measurement 03-34-08PM~ |
| Time Captured: | 15:34:38 27/02/2020 |

**Capture Settings**

| | |
|---|---|
| Camera Type: | sCMOS |
| Laser Type: | Blue488 |
| Camera Level: | 16 |
| Slider Shutter: | 1300 |
| Slider Gain: | 512 |
| FPS | 25.0 |
| Number of Frames: | 2248 |
| Temperature: | 25.0 - 25.0 °C |
| Viscosity: | (Water) 0.888 - 0.889 cP |
| Dilution factor: | 2 x 10e3 |
| Syringe Pump Speed: | 70 |

**Analysis Settings**

| | |
|---|---|
| Detect Threshold: | 5 |
| Blur Size: | Auto |
| Max Jump Distance: | Auto: 12.7 - 14.4 pix |

**Results**

| Stats: Merged Data | |
|---|---|
| Mean: | 137.1 nm |
| Mode: | 96.1 nm |
| SD: | 76.1 nm |
| D10: | 72.2 nm |
| D50: | 116.9 nm |
| D90: | 224.6 nm |

| Stats: Mean +/- Standard Error | |
|---|---|
| Mean: | 136.0 +/- 4.8 nm |
| Mode: | 99.9 +/- 4.4 nm |
| SD: | 73.1 +/- 8.4 nm |
| D10: | 72.0 +/- 1.4 nm |
| D50: | 116.4 +/- 4.0 nm |
| D90: | 223.7 +/- 5.4 nm |
| Concentration (Upgrade): | 2.78e+11 +/- 3.24e+10 particles/ml |
| | 22.0 +/- 3.0 particles/frame |
| | 27.8 +/- 4.4 centres/frame |

**(B)**

**Details**

| | |
|---|---|
| NTA Version: | NTA 3.4 - Sample Assistant Build 3.4.003-SA |
| Script Used: | SOP AutoSampler Measurement 03-34-08PM~ |
| Time Captured: | 15:34:38 27/02/2020 |

**Capture Settings**

| | |
|---|---|
| Camera Type: | sCMOS |
| Laser Type: | Blue488 |
| Camera Level: | 16 |
| Slider Shutter: | 1300 |
| Slider Gain: | 512 |
| FPS | 25.0 |
| Number of Frames: | 2248 |
| Temperature: | 25.0 - 25.0 °C |
| Viscosity: | (Water) 0.9 cP |
| Dilution factor: | 5 x 10e2 |
| Syringe Pump Speed: | 70 |

**Analysis Settings**

| | |
|---|---|
| Detect Threshold: | 5 |
| Blur Size: | Auto |
| Max Jump Distance: | Auto: 14.5 pix |

**Results**

| Stats: Merged Data | |
|---|---|
| Mean: | 92.5 nm |
| Mode: | 68.5 nm |
| SD: | 44.1 nm |
| D10: | 48.7 nm |
| D50: | 80.8 nm |
| D90: | 152.7 nm |

| Stats: Mean +/- Standard Error | |
|---|---|
| Mean: | 92.0 +/- 6.1 nm |
| Mode: | 70.7 +/- 3.5 nm |
| SD: | 42.6 +/- 3.6 nm |
| D10: | 49.7 +/- 3.6 nm |
| D50: | 81.4 +/- 5.5 nm |
| D90: | 153.3 +/- 17.7 nm |
| Concentration (Upgrade): | 9.66e+09 +/- 6.36e+08 particles/ml |
| | 2.9 +/- 0.2 particles/frame |
| | 4.3 +/- 0.3 centres/frame |

**Supplementary Figure 2. Malvern NS300 NTA settings**. The settings recorded above were used to analyse **(A)** ultracentrifugation **(B)** tangential flow filtration isolated HEK293 EVs to determine batch EV size and concentration. Settings and data shown are of representative samples shown in Figures 1E & 1F.

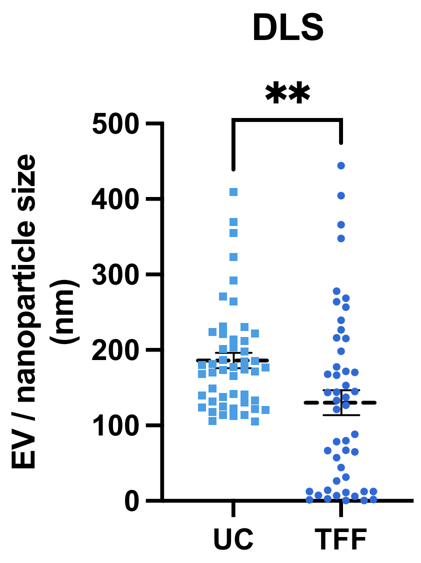


**Supplementary Figure 3. Dynamic light scattering size analysis of ultracentrifugation (UC) and tangential flow filtration (TFF) isolated HEK293 EV batches.** Error bars denote standard error of the mean, $n$ = 49 EV batches, student $t$-test **$P$<0.01.

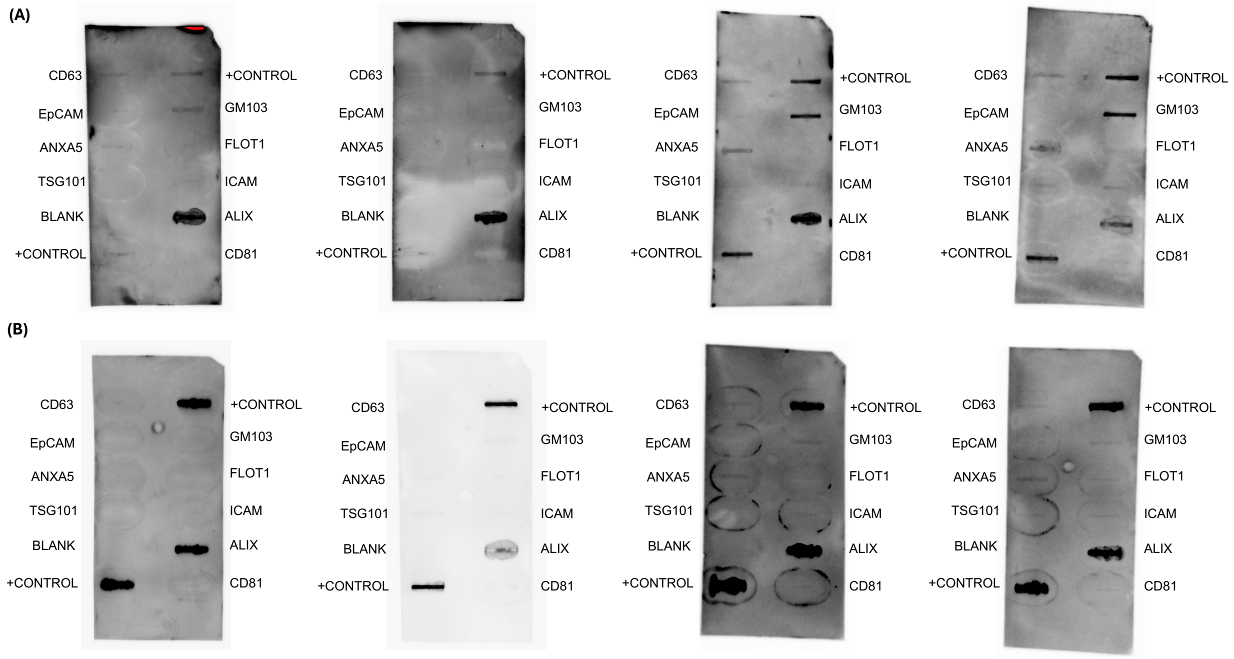


**Supplementary Figure 4. Exo-Check Exosome Antibody Array analysis of HEK293 cell-derived extracellular vesicles.** Four independent **(A)** ultracentrifugation (UC) and **(B)** tangential flow filtration (TFF) isolated HEK293 EVs were processed and analysed for the presence of EV markers using the Exo-Check Exosome Antibody Array Kit. The Exo-Check array included 12 dot blot spots/lines that incorporated antibodies specific for the EV markers CD63, CD81, ALIX, FLOT1, ICAM, EpCam, ANXA5 and TSG101, as well as a cellular contamination control (GM130 cis-Golgi marker) and several assay positive [+control] and negative [blank] controls. Developed dot blot arrays were imaged using a Bio-Rad ChemiDoc imaging system. Two images were selected from this panel for inclusion in Figures 2A & 2B respectively.

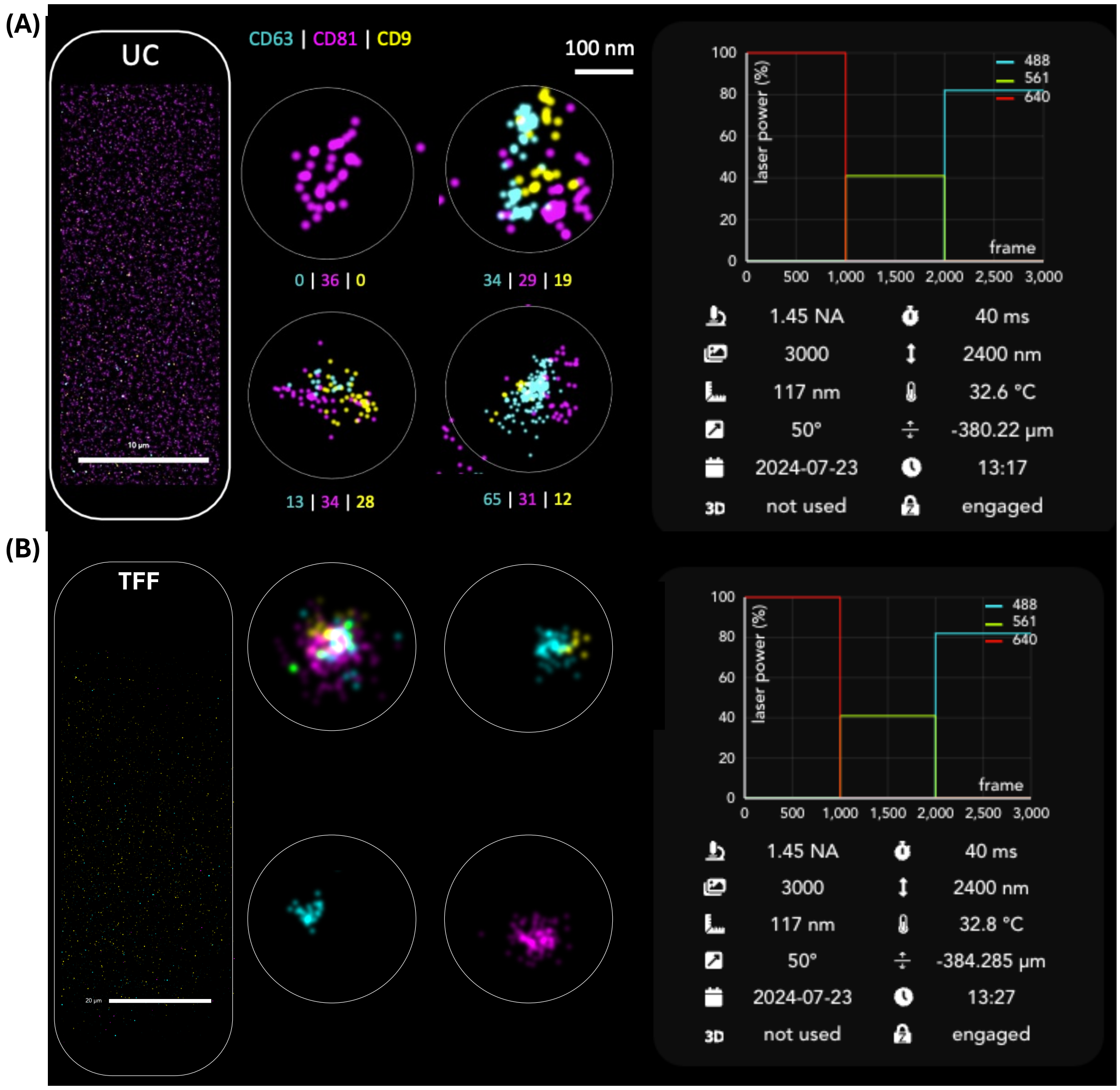


**Supplementary Figure 5 Super resolution microscopy of HEK293 extracellular vesicles.** Representative super resolution images of **(A)** Ultracentrifugation (UC) isolated extracellular vesicles and **(B)** tangential flow filtration (TFF) isolated extracellular vesicles. Settings shown above including 1.45 numerical aperture (NA), 3000 frames, 50° illumination angle, 40 ms exposure time, 2400 nm axial distance from focus reference, 32.6 °C or 32.8°C temperature, -380.22 µm or -384.285 µm focus value with Z-lock engaged.

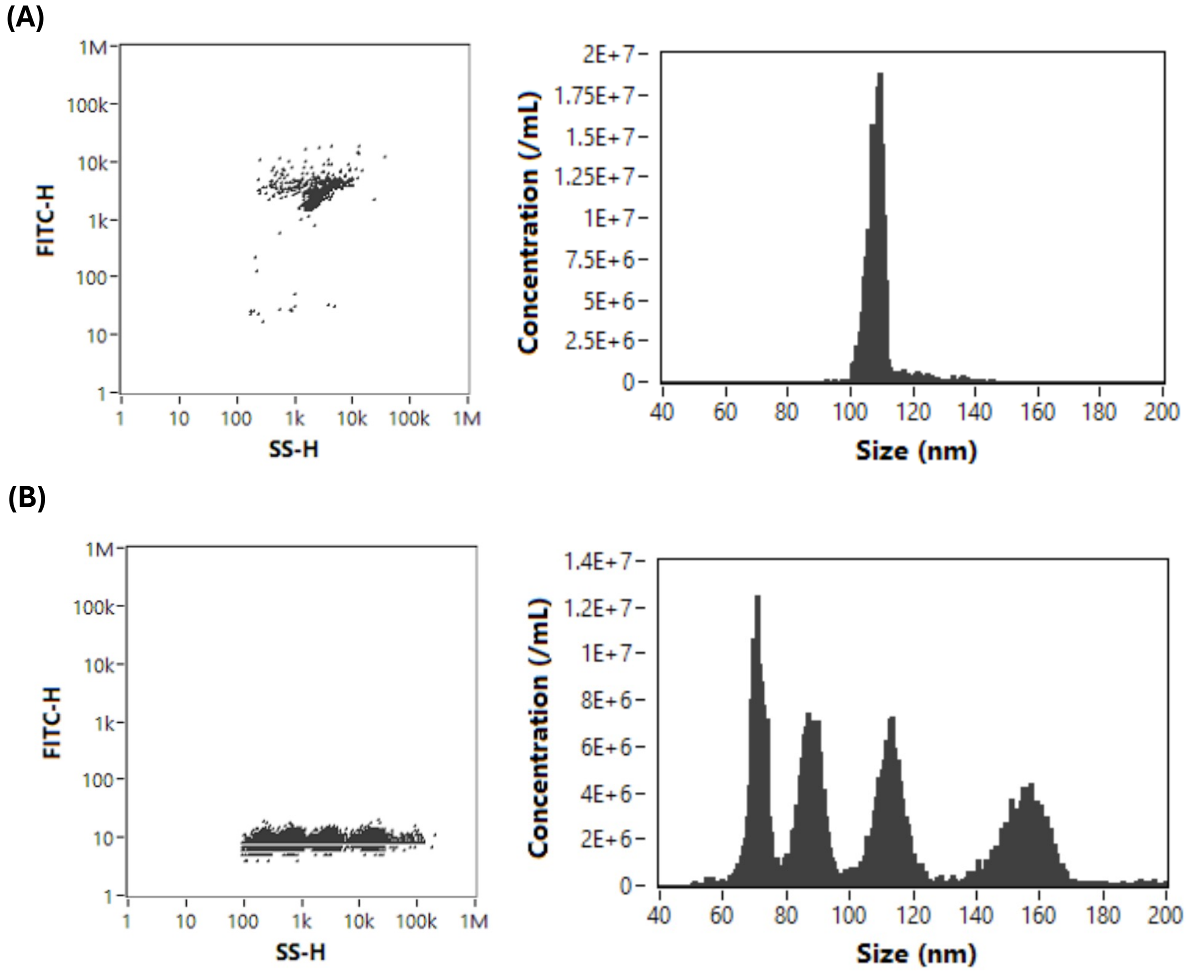


**Supplementary Figure 6. Nano flow cytometry QC and size calibration beads.** Representative nano flow cytometry scatter plots and size histograms of **(A)** 250 nm SiNP Dual laser QC Beads (2.16E+10; #QS2503; NanoFCM, China) and **(B)** size reference standard silica nanospheres (68–155 nm; #S16M-Exo; NanoFCM, China).

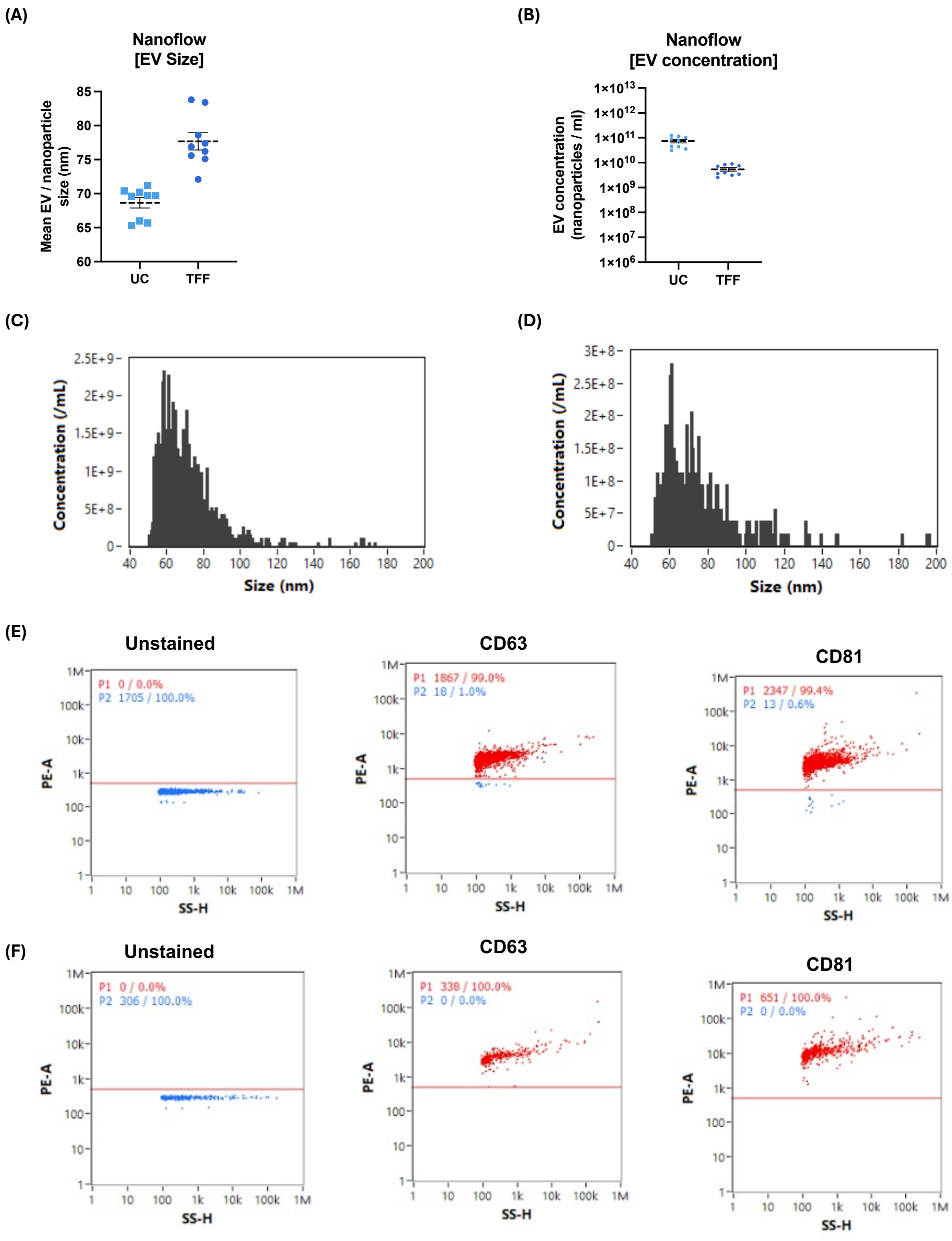


**Supplementary Figure 7. Nanoflow cytometry characterisation of CD63 and CD81 positive HEK293 extracellular vesicles (EVs).** Mean EV/nanoparticle **(A)** size and **(B)** concentration of ultracentrifugation (UC) and tangential flow filtration (TFF)-isolated HEK293 EVs that were either unstained or stained with CD63-PE or CD81-PE antibodies. These data relate to Figure 2C in the main manuscript. Error bars denote standard error of the mean. Representative size histograms of **(C)** UC and **(D)** TFF-isolated EVs and representative scatter plots including gating strategy of unstained, CD63-PE and CD81-PE antibody stained **(E)** UC and **(F)** TFF-isolated HEK293 EVs are also shown as indicated.

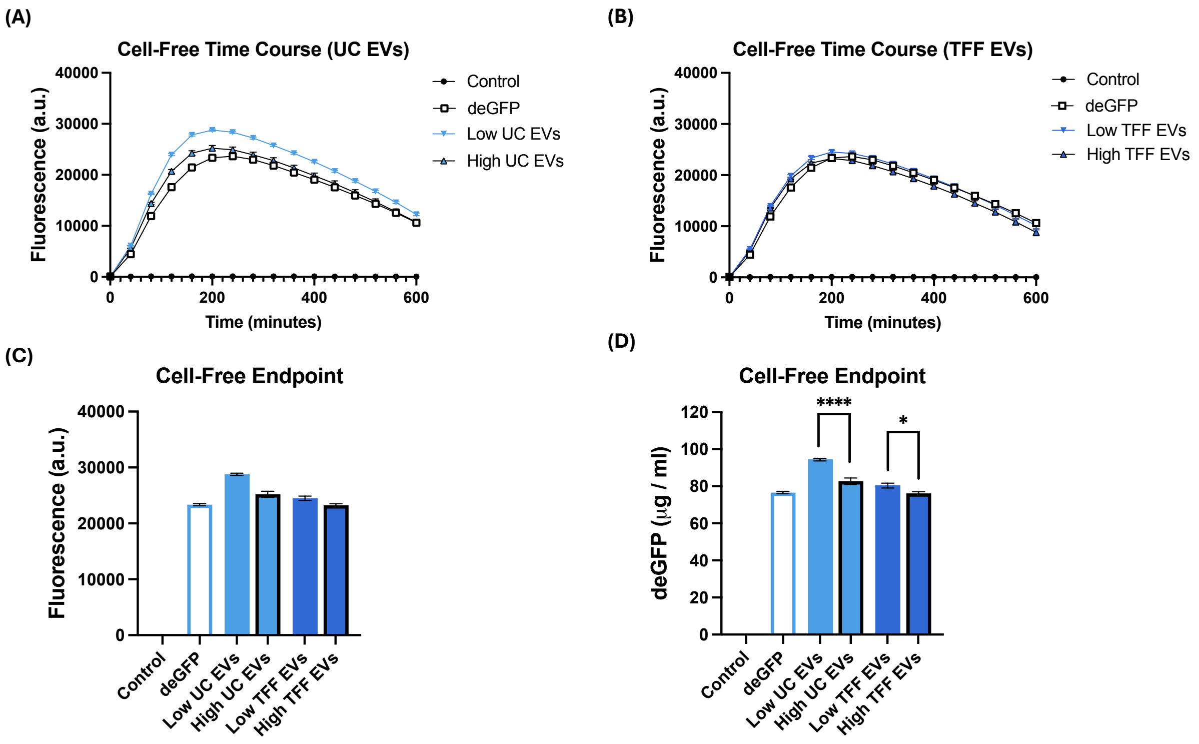


**Supplementary Figure 8. Impact of extracellular vesicle (EV) concentration on cell-free reaction performance.** Time course analysis of 10 µL-scale cell-free reactions expressing deGFP in the presence of 0 µl, 0.3 µl [Low] or 0.6 µl [High] concentrations of **(A)** UC or **(B)** TFF-isolated HEK293 cell EVs ($2.5 \times 10^{10}$ / ml) as indicated. Control denotes cell-free reactions containing 0 µl EVs and 0 ng deGFP plasmid, whilst deGFP denotes cell-free reactions containing 0 µl EVs and 400 ng deGFP plasmid. Endpoint (200 minutes) **(C)** fluorescence and **(D)** GFP curve calibrated measurements of the indicated cell-free reactions are also shown. deGFP fluorescence was measured using a BMG CLARIOstar plate reader (excitation 483-14 nm, 502.5 nm dichroic, emission 530–30 nm and 1000 gain). Error bars denote standard error of the mean, $n = 9$ independent cell-free reactions, student $t$-test $*P<0.05$, $****P<0.0001$.

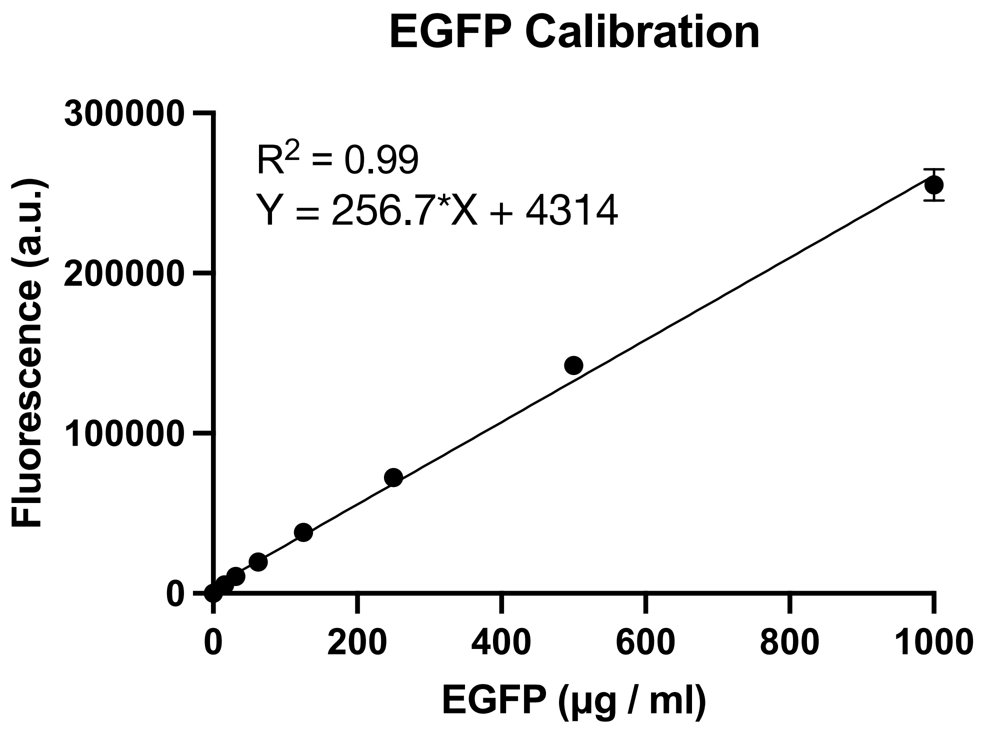


**Supplementary Figure 9. EGFP calibration curve.** Error bars denote standard error of the mean, n=4 technical measurements.

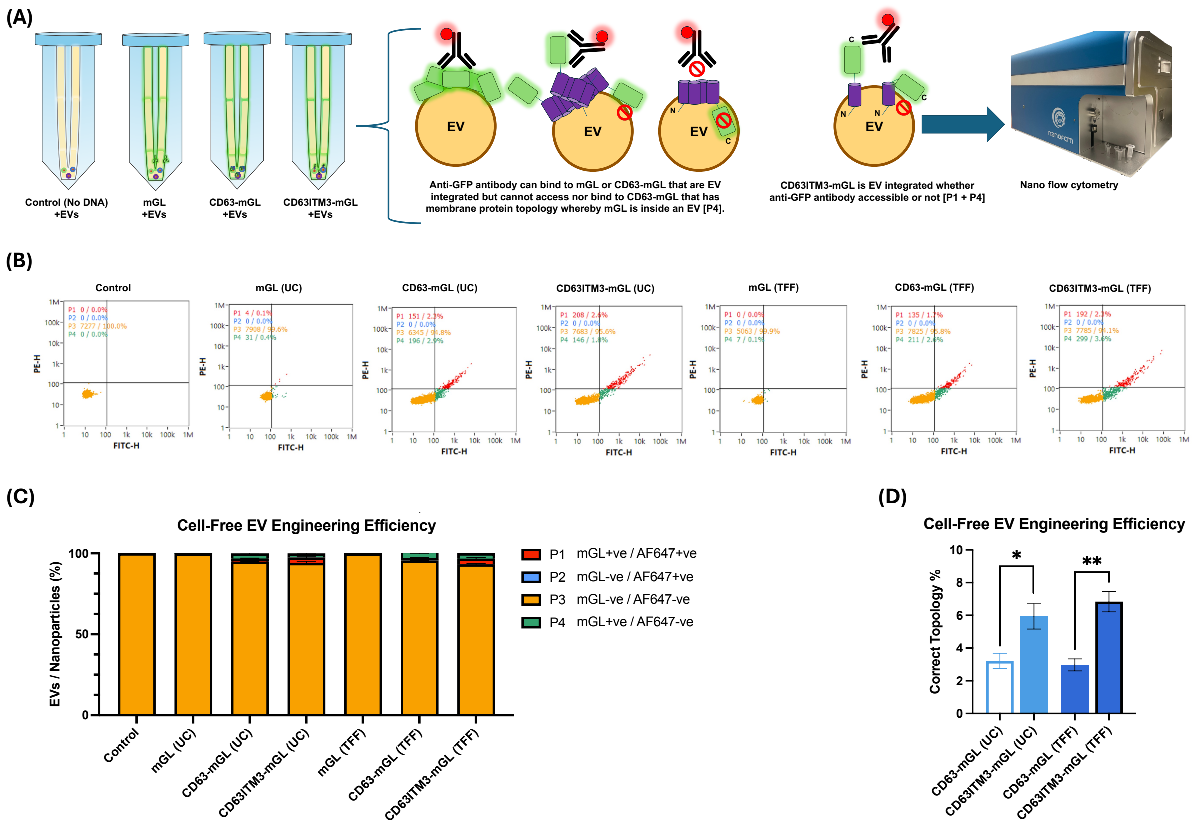


**Supplementary Figure 10. Cell-free extracellular vesicle engineering efficiency. (A)** Assay workflow for cell-free EV engineering and nano flow cytometry analysis of ultracentrifugation (UC) and tangential flow filtration (TFF)-isolated HEK293 EVs engineered in control (no DNA), mGL, CD63-mGL or CD63ITM3 cell-free reactions as indicated. **(B)** Representative nano flow cytometry scatter plots of UC or TFF-isolated HEK293 EVs that were cell-free engineered in control or protein producing reactions (mGL, CD63-mGL or CD63ITM3-mGL) and then stained with anti-GFP Alexa Fluor (AF)-647 antibody prior to analysis. A quadrant gating strategy (P1-P4) was used to identify nanoparticle/EV subpopulations based upon their FITC-H (mGL, CD63-mGL or CD63ITM3-mGL detection) and PE-H (670/30 nm; anti-GFP AF-647 antibody) detection signals. **(C)** Percentage (%) of total analysed nanoparticles/EVs from cell-free EV engineering reactions within each of the gated quadrants P1 (FITC + / + Anti-GFP AF-647), P2 (FITC - / + Anti-GFP AF-647), P3 (FITC - / - Anti-GFP AF-647) and P4 (FITC + / - Anti-GFP AF-647). **(D)** Percentages of UC and TFF-isolated EVs that were successfully cell-free EV engineered with CD63-mGL (P4) or CD63ITM3-mGL (P1+P4). Error bars denote standard error of the mean, $n = 3$ independent cell-free reactions, student *t*-test $*P<0.05$, $**P<0.01$.

**Supplementary table 1. Protein sequences**

| Protein & description | Sequence | Reference(s) |
|---|---|---|
| **deGFP**<br><br>**eGFP-Del6-229**<br><br>(Enhanced green fluorescent protein [EGFP] variant) | MELFTGVVPILVELDGDVNGHKFSVSGEGEGDATYGKLTL KFICTTGKLPVPWPTLVTTLTYGVQCFSRYPDHMKQHDFF KSAMPEGYVQERTIFFKDDGNYKTRAEVKFEGDTLVNRIE LKGIDFKEDGNILGHKLEYNYNSHNVYIMADKQKNGIKV NFKIRHNIEDGSVQLADHYQQNTPIGDGPVLLPDNHYLST QSALSKDPNEKRDHMVLLEFVTAAGI | [1] & This study |
| **mGL**<br><br>monomeric green lantern (Green fluorescent protein [GFP] variant) | MVSKGEELFTGVVPILVELDGDVNGHKFSVRGEGEGDAT NGKLTLKFICTTGKLPVPWPTLVTTLGYGVACFARYPDHM KQHDFFKSAMPEGYVQERTISFKDDGTYKTRAEVKFEGDT LVNRIVLKGIDFKEDGNILGHKLEYNFNSHKVYITADKQK NGIKANFKTRHNVEDGGVQLADHYQQNTPIGDGPVLLPD NHYLSHQSKLSKDPNEKRDHMVLKERVTAAGITHDMDEL YK | [2] & This study |
| **CD63-mGL**<br><br>CD63 tetraspanin protein fused with a 20 amino acid linker and mGL. | MAVEGGMKCVKFLLYVLLLAFCACAVGLIAVGVGAQLV LSQTIIQGATPGSLLPVVIIAVGVFLFLVAFVGCCGACKENY CLMITFAIFLSLIMLVEVAAAIAGYVFRDKVMSEFNNNFRQ QMENYPKNNHTASILDRMQADFKCCGAANYTDWEKIPSM SKNRVPDSCCINVTVGCGINFNEKAIHKEGCVEKIGGWLR KNVLVVAAAALGIAFVEVLGIVFACCLVKSIRSGYEVMGG QSGGSQGQSSGQQSSQSSMVSKGEELFTGVVPILVELDGD VNGHKFSVRGEGEGDATNGKLTLKFICTTGKLPVPWPTLV TTLGYGVACFARYPDHMKQHDFFKSAMPEGYVQERTISF KDDGTYKTRAEVKFEGDTLVNRIVLKGIDFKEDGNILGHK LEYNFNSHKVYITADKQKNGIKANFKTRHNVEDGGVQLA DHYQQNTPIGDGPVLLPDNHYLSHQSKLSKDPNEKRDHM VLKERVTAAGITHDMDELYK | UniProt [CD63; P08962] & This study |
| **CD63ITM3-mGL**<br><br>CD63 transmembrane domain 3 fused with a 20 amino acid linker and mGL | MYCLMITFAIFLSLIMLVEVAAAIAGYVFRDKVMSEFNNN FRQQMENYPKNNHTAGGQSGGSQGQSSGQQSSQSSMVSK GEELFTGVVPILVELDGDVNGHKFSVRGEGEGDATNGKLT LKFICTTGKLPVPWPTLVTTLGYGVACFARYPDHMKQHDF FKSAMPEGYVQERTISFKDDGTYKTRAEVKFEGDTLVNRI VLKGIDFKEDGNILGHKLEYNFNSHKVYITADKQKNGIKA NFKTRHNVEDGGVQLADHYQQNTPIGDGPVLLPDNHYLS HQSKLSKDPNEKRDHMVLKERVTAAGITHDMDELYK | [3] & This study |

**Supplementary references**


1. Shin, J. and Noireaux, V. (2010) Efficient cell-free expression with the endogenous E. Coli RNA polymerase and sigma factor 70. *J Biol Eng* 4, 8
2. Campbell, B.C. *et al.* (2020) mGreenLantern: a bright monomeric fluorescent protein with rapid expression and cell filling properties for neuronal imaging. *Proceedings of the National Academy of Sciences* 117, 30710–30721
3. Curley, N. *et al.* (2020) Sequential deletion of CD63 identifies topologically distinct scaffolds for surface engineering of exosomes in living human cells. *Nanoscale* 12, 12014–12026
